# From empirical to data-driven host selection: a broad-host-range expression platform to facilitate chassis screening

**DOI:** 10.1101/2022.08.09.503317

**Authors:** Viviënne Mol, Kristoffer Bach Falkenberg, Ácil De Almeida Will, Ivan Pogrebnyakov, Charlotte Beck, Anna Lyhne Skøttrup, Alex Toftgaard Nielsen, Sheila Ingemann Jensen

**Author notes:** Correspondence to: Sheila Ingemann Jensen, The Novo Nordisk Foundation Center for Biosustainability, Technical University of Denmark, 2800 Kgs. Lyngby, Denmark.

## Abstract

Nature has provided a vast landscape of organisms through evolution, each with unique phenotypic traits adapted to varying environments. Nevertheless, host selection in biotechnological research is exceedingly dominated by empirical preference, where the endogenous physiology of the selected host is often not suited to the desired application. Considering that large parts of cellular regulation and metabolism remain obscure, empirical selection of a preferred model organism may lead to undue caveats in further engineering attempts, arising from intrinsic metabolism. One reason for the empirical host selection is the lack of engineering tools for screening novel organisms. In this study, we provide a modular, single vector-based expression platform, compatible with a wide range of prokaryotes. It centers around a tight and titratable promoter system, inducible by anhydrotetracyclin within an 84-fold dynamic range. It enables easy screening of recombinant proteins and pathways in both mesophilic and thermophilic Gram-negative and Gram-positive hosts. Overall, this platform enables simple screening of heterologous expression and production in a variety of hosts, including the exploration of previously unconsidered hosts thereby aiding the transition from empirical to data-driven host selection.

## Introduction

In the past decade, there has been an unprecedented increase in synthetic biology applications in all areas of biotechnology. This has facilitated an improved understanding and treatment of disease, as well as providing critical tools for the generation of cell factories for mitigating climate change through the transition into a circular bio-economy and the production of alternative proteins. Generally, most studies involving synthetic biology rely on the use of model organisms due to their genetic accessibility through the available molecular tools and the relatively thorough understanding of genotypes and phenotypes, compared to other hosts. For health and disease, the use of model organisms is critical to subsequently translate findings into human systems. However, although this consideration is less critical when developing microbial bioproduction solutions, a heavy reliance on a subset of model organisms still dominates the field.

Since 1884, prokaryotes have been classified according to their response to Gram staining, resulting from the fundamental differences found in the cell envelope structure of the organism^1^. Throughout time, though further isolation and identification of a vast range of microbial phenotypes occurred, history dictated a predominant use of specific model microbial hosts such as the Gram-negative *Escherichia coli*^2,3^ and Gram-positive *Bacillus subtilis*^4–6^ in biotechnology. As a result, together with challenges in the successful isolation of environmental isolates, a large fraction of the microbial metabolic potential lies under-studied and underdeveloped, forming a huge untapped resource to tackle some of society’s most pressing issues. Combining this metabolic potential with the premise of synthetic biology relies on tools that facilitate easy screening of functionality across a range of microorganisms. Though there is an increased interest in using novel microbes for bioproduction purposes, tools developed are predominantly targeted to the specific unique host^7^. The often empiric and preferential host selection for bioproduction purposes results in working in silos, and both academia and industry could therefore greatly benefit from more broad-host range applicable tools. This can ameliorate the need to constantly develop organism specific tools, at least for the initial stages of research.

Through the development of synthetic biology, significant knowledge has been gathered about metabolism and genotype-phenotype relationships, but due to the complex nature of life, there is still a lack of ability to rationally engineer specific complex traits, such as tolerance or thermostability. Thus, relying on the engineering of favored model organisms to introduce these traits is not trivial. Fortunately, through evolution, nature has created vast landscapes of phenotypically diverse microorganisms, each with tailored, adaptive metabolisms fit to specific conditions and objectives. Thus, the selection of a host that has a fitting metabolic profile or protein production machinery for a process or product of interest is favored. With the constant development of synthetic biology tools, the ability to harness and engineer this vast landscape increases, providing immense novel bioproduction potential. To that end, letting host metabolism guide host selection, instead of a biased predisposition to a host based on empirical preference, can facilitate the application of microbial cell factories to favor the green transition that is so critically needed. Though acknowledged across research institutions and industries, this is not as trivial as it may seem^7,8^. To highlight this, the Gram-negative *Pseudomonas putida* is often perceived to show superior tolerance to various cellular stresses. However, comparative toxicity studies between *P. putida* and *E. coli* show that the former is not alwas the more tolerant organism, highlighting that toxicity is hard to predict and is dependent on both the compound and the species^9^. This demonstrates the drawbacks of certain *a priori* assumptions and emphasizes the need to evaluate on a case-by-case basis. Additionally, the majority of proteins purified for crystallographic purposes still rely on significant engineering and optimization in *E. coli* production hosts, which could be avoided if more favorable expression hosts are used^10,11^. Aside from microbial bioproduction and protein crystallographic purposes, fundamental studies can also significantly benefit from studying protein function across hosts. As a protein may fold and hence function differently across contexts, dependent for example on complex formation, studies across hosts can elucidate additional information about its structure and function^12^. However, predominantly in laboratories where experience working with different organisms is lacking, the laborious molecular biology needed to create vectors for various hosts presents a significant barrier. Additionally, when novel biotechnological inventions are patented, for broad claims, functionality across the phylogenetic space should be demonstrated. Therefore, to bridge this gap broad-host-range expression vectors are required.

Whereas most of the studies that have focused on developing broad-host-range vectors concentrate on identifying replication and resistance cassettes^13,14^, one critical aspect that is often unconsidered and hence underdeveloped is a broad-host-range inducible expression system. Research has shown that the efficiency of expression systems can vary greatly between hosts. As an example, developed expression systems can lose tightness and titratability between the (relatively related) *E. coli* and *P. putida*, ^15^. Ideally, a broad-host-range expression system functions similarly across a wide variety of hosts, with specific characteristics being critical for functioning. Inducible systems are often desired to provide control over the timing of expression and facilitate the expression of toxic or metabolically burdening proteins. This comes with the added requirement of being silent in the uninduced state. As an added desired trait, linear titratability across a defined range is beneficial, as the flexibility of lowered expression can prove to be beneficial for protein translation levels and hence protein folding^16^. Finally, homogenous expression across the population is instrumental for accurate strain analysis and comparison, as well as to avoid non-productive sub-populations in production processes.

Aside from functional characteristics, to obtain a broad-host-range cassette, the DNA sequence must be compatible with a range of different hosts. One critical aspect that is often overlooked when working with different host organisms is the restriction-modification system, a defense mechanism preventing the maintenance of foreign DNA, found across the prokaryotic kingdom^17^. Though difficult to predict, preventing the presence of certain recognition sites can favor the transformation of a foreign genetic construct into various hosts.

As cells have evolved to respond to complex biological inputs, there is a vast variety of inducible expression systems found in nature, needed to provide dynamic genetic regulation. One inducible expression system is based upon the Tet repressor (TetR) protein, in combination with the *tet* operator, naturally found in *E. coli* and involved in controlling tetracycline resistance^18,19^. In *E. coli* it has been shown to possess the desired characteristics mentioned above (tight, titratable and homogeneous) when induced with a tetracycline analog, anhydrotetracycline (aTc)^19^. To function in *B. subtilis*, the *tet* regulatory elements were inserted and modified in the strong xylose promoter (P_xyl_)^20,21^. This variant was previously shown to be silent, titratable and homogenous in *B. subtilis*, as well as silent in an *E. coli* DH5α cloning strain, critical to allowing easy cloning of desired target genes^22^. Different variants of this promoter have also been shown to function in *Actinomyces oris*^23^, *Staphylococcus* species^24–26^, *Bacillus licheniformis*^27^, *Clostridium difficile*^28,29^, *Chlamydia trachosomatis*^30^, and *Acetobacterium woodii*^31^, however systematic analysis of whether one variant can function across hosts is lacking. Additionally, the differences between variants used make the comparison across hosts difficult. Yet this highlights the potential of the TetR-P_tet_ system across prokaryotic phyla, and hence the TetR-P_tet_ system was used as a starting point to build a broad-host-range expression vector that can be used across a range of prokaryotic organisms, including Gram-positive, Gram-negative and both mesophilic and thermophilic hosts.

In this study, we provide a modified TetR-P_tet_ expression system that allows screening of proteins and pathways across various relevant prokaryotes. To show the applicability of the system, we demonstrate screening various enzymes for the production of aromatic phenolic compounds across three distinct microbial production hosts, using a single plasmid construct. The availability of this system can allow more systematic studies across hosts, both for bioproduction as well as fundamental purposes, and hence facilitate more data-driven selection of hosts.

## Results

### Modification of the TetR-P_tet_ sequence allows broad-host-range functionality

To investigate the applicability of the TetR-P_tet_ promoter as a broad-host-range tight inducible expression system, we selected four species as test-bed for broad-host-range functionality in this study; *E. coli, P. putida, B. subtilis* and *Parageobacillus thermoglucosidasius*. Classically *E. coli* is a Gram-negative organism that has extensively been used for fundamental studies and as a production workhorse, owing to its long history of use in the field of molecular biology^2,3^. *P. putida* has gained interest as a microbial host suitable for cell factory engineering. While *P. putida* is also a Gram-negative host, it has a different substrate metabolism and tolerance to oxidative stress^32^. Though metabolically different, *P. putida* has a phylogenetic origin relatively close to that of *E. coli*, allowing the application of various engineering and synthetic biology tools to be translated between organisms, albeit with some necessary modifications^15^.

The absence of a second outer phospholipid membrane makes Gram-positive organisms the predominant choice for industrial protein secretion^4–6^. *B. subtilis* is considered a model Gram-positive organism for protein production and secretion. Species in the *Bacillus* genus have been used across industries as the largest producers of industrial enzymes to date^33^. Though some *Bacilli* species have a flexible growth temperature range up to 50 °C^34^, they are still considered mesophilic hosts. For the production of valuable biochemicals, an increased processing temperature has various advantages such as improved substrate and product solubility, lowered viscosity and improved mass transfer kinetics^35^. Additionally, for volatile products, *in situ* product removal is facilitated and beneficial to simplify downstream processing costs, as well as prevent product toxicity and pathway inhibition to production strains^36^. For thermophilic processes, *P. thermoglucosidasius* is a valuable host with a wide substrate utilization profile and proven production potential^36,37^.

Where the TetR-P_tet_ expression system has often been used in its native host, *E. coli*, significant modifications to the promoter were needed to allow efficient functioning in other hosts. Initially, the endogenous *tet* regulatory elements from *E. coli* (Figure 1A) were combined with the strong *B. subtilis* xylose (P_xyl_) promoter, with modified −35 and −10 elements to allow the functioning of the promoter in the Gram-positive *B. subtilis*^20^. Since then, for use in other hosts such as *C. difficile*^29^ and *A. woodii*^31^, slight modifications have been made, targeted to the specific host. To our knowledge, all studies using variants of the tetracycline inducible system only show application within one bacterial phylum. Here, the sequence as previously proposed in *C. difficile*^28^ and thoroughly characterized in *B. subtilis*^22^, was used as starting sequence.

**Figure 1:**
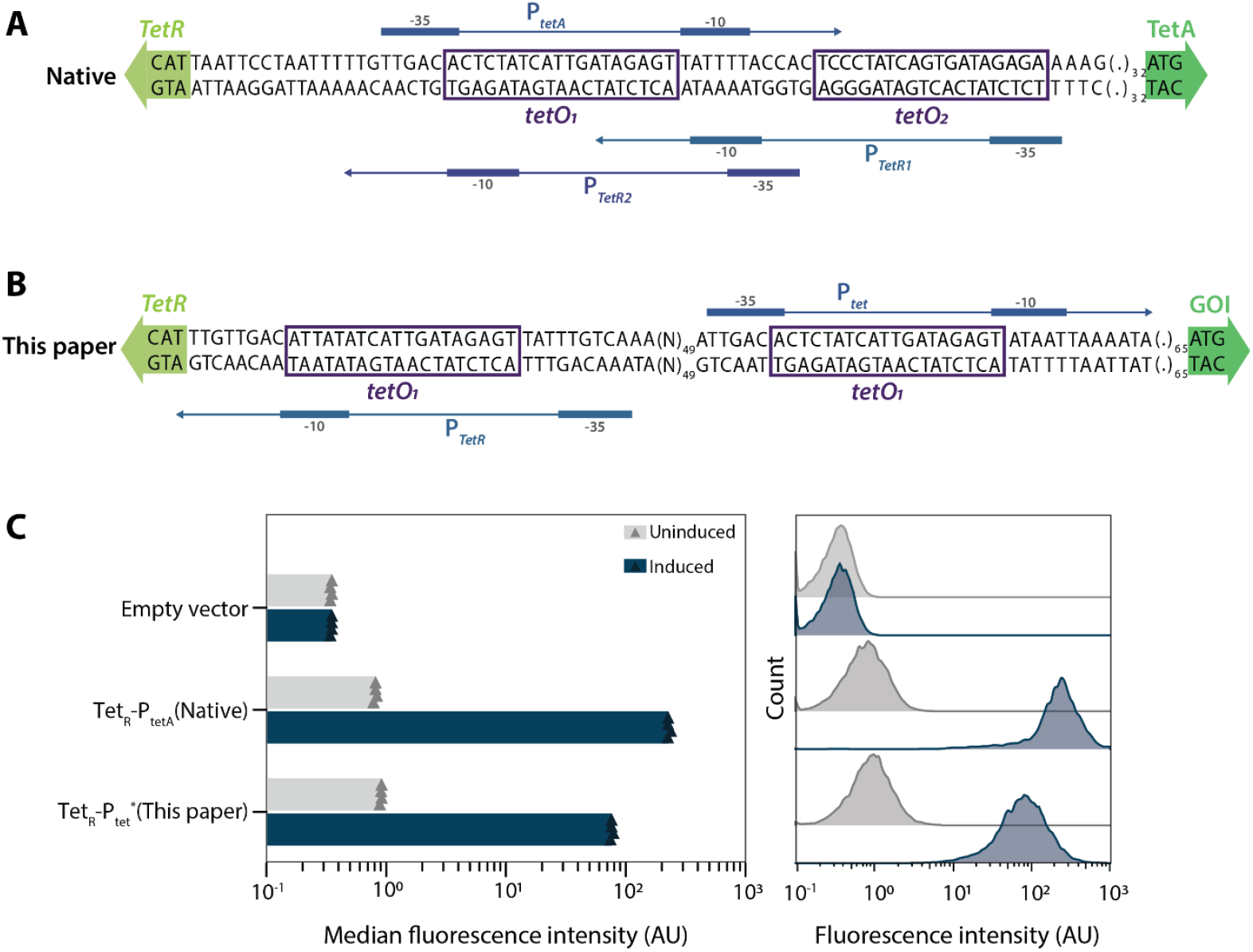
Modification of the TetR-P_tetA_ promoter in *E. coli*. The organization of the native *E. coli* TetR-P_tetA_ promoter (A)^18^ and the promoter as used in this paper, TetR-P_tet_^*^ (B) are shown. The start of coding sequences of the TetR protein and a possible Gene of Interest (GOI) are represented by green arrows. Purple boxes indicate the *tetO* sites. For the P_tetA_, P_TetR1_ and P_TetR2_ promoters, the −35/−10 and transcription start sites are shown as thicker lines and arrowheads, respectively. (.)_X_ represents the 5’ untranslated region, composed of X bases. (N)_X_ represents a spacer sequence of X basepairs long between the P_TetR_ and P_tet_. Characterization of *E. coli* cultures expressing either TetR-P_tet_ system controlling mRuby2 or an empty vector (pSEVA631), 8 hours post induction with 1.0 µg mL^−1^ aTc (C). Population data from a representative sample of each condition is shown, where biological quadruplicates are summarized according to the mean of individual median fluorescence intensity, with replicate medians shown as triangles.

For use in other hosts, the presence of restriction modification (RM) systems can interfere with construct maintenance and so recognition of the construct by RM systems should be avoided^38^. Therefore, we searched the TetR-P_tet_ sequence for restriction enzyme (RE) recognition sites that are unique to the TetR-P_tet_ sequence when compared to other sequences known to transform into a range of hosts (for example the pBBR1 and pAMβ1 sequences). In the original TetR-P_tet_ sequence, seven unique sites were identified (Table 1). Many prokaryotic genomes encode restriction enzymes, yet predicting what sequence they may recognize is currently impossible. Additionally, the REs corresponding to the unique recognition sites found have been identified across several phyla, and so for application across the prokaryotic phylogenetic tree, all seven RE recognition sites were removed from the TetR-P_tet_ sequence. Nine silent mutations were introduced in the TetR coding sequence, and four other A→T or G→C mutations were introduced into the 5’ untranslated region of the P_tet_ promoter (Figure 1B, Supplementary sequences). Where transformation of a vector containing the TetR-P_tet_ sequence with RE sites yielded no colonies upon transformation into *P. thermoglucosidasius*, we observed improved transformation efficiencies of the same vector with the RE sites removed (from here on designated as TetR-P_tet_^*^).

**Table 1:** Restriction enzyme recognition sites with corresponding restriction enzymes and isochimers, which were unique in the TetR-P_tet_ sequence compared to DNA sequences known to be transformable in a range of different microbes. Organisms from which the restriction enzymes were isolated are indicated.

| Recognition sequence | Restriction enzyme | Identified in |
| --- | --- | --- |
| ACNNNNNCTCC | BsaXI | *Bacillus stearothermophilus* 25B |
| CACNNNGTG | AleI, OIiI | *Aureobacterium liquefaciens*, *Oceanospirillum linum* 4-5D1 |
| CCCACA | RleAI | *Rhizobium leguminosarum* DSM 5629 |
| CCTNNNNNAGG | BstENI, EcoNI, XagI | *Geobacillus stearothermophilus* EN, *Escherichia coli* CDC A-193 (ATCC 12041), *Xanthobacter agilis* Vs18-132 |
| GAGCTC | SacI, Eco53kI, BpuAml, Ecl136II, EcoICRI, MxaI, Psp124BI, SstI | *Streptomyces achromogenes* (ATCC 12767), *Escherichia coli* 53k, *Bacillus pumilus*, *Enterobacter cloacae* RFL136, *Myxococcus xanthus* F18E, *Pseudomonas* sp. 124B, *Streptomyces stanford* |
| GTTAAC | BstEZ359I, BstHPI, HpaI, KspAI, SsrI | *Bacillus stearothermophilus* EZ359, *Bacillus stearothermophilus* HP, *Haemophilus parainfluenzae*, *Kurthia* sp. N88, *Staphylococcus saprophyticus* B6 |
| TACGTA | BstSNI, Eco105I, SnaBI | *Bacillus stearothermophilus* SN, *Escherichia coli* RFL105, *Sphaerotilus natans* |

Aside from modifications to the promoter that allow broad-host-range transcription, efficient translation across species was also considered. Translation efficiency is dependent on various sequence elements, such as the Shine-Dalgarno (SD) sequence and its spacing to the start codon, which are targets for optimizing protein expression in specific hosts^39–41^. Whole genome sequencing and transcriptomic data can provide insight into consensus sequences across species^42,43^, though, to our knowledge, no experimental comparative studies of consensus sequences across hosts have been performed to date. Most SD sequences contain slight variations of the conserved AGGAGG core sequence^40,41,43,44^, and hence this was selected as a broad-host-range SD sequence for further work. The spacer between the SD and start codon is known to be less stringent in *E. coli* and varies from 5 to 13 nucleotides^41,42^. Comparative analysis of 18 genomes has shown a consensus spacer from 5 to 10 nucleotides in length^45^, matching reports on suitable spacer lengths specific to various hosts^41,44,46,47^. Considering *Bacilli* have been shown to have more stringent spacer requirements, a broad-host-range spacer of 7 nucleotides in length was selected^44^.

Comparison of the native *E. coli* TetR-P_tet_A sequence versus the modified, broad-host-range TetR-P_tet_^*^ sequence was done through analysis of mRuby2 expression^48^ (Figure 1C). Single cell characterization showed that removing one *tet* operator (tetO) sequence does not come at a cost of tightness when uninduced or of homogeneity when induced. The TetR-P_tet_^*^ promoter remained highly inducible, though with some loss of its dynamic range, but still maintaining an 84-fold change in mRuby2 expression upon induction. Both variants are stable up to at least 8 hours post induction, with the speed of response unchanged (Supplementary Figure S1). The desired characteristics verified in the *E. coli* host enables further validation of the expression system across other bacterial phyla.

### TetR-P_tet_^*^ is a favorable broad-host-range expression system

Throughout literature, various examples of the application of the TetR-P_tet_ system can be found (Figure 2A). However, between these examples, different variants of the TetR-P_tet_ sequence are used. For example, work done in *P. putida*^49^ and two M*ycobacterium*^50^ strains was performed using the native *E. coli* system, though this version does not function in *B. subtilis*^20^. Therefore, generally for each host slight modifications have been introduced, both intentionally and incoherently. To screen multiple hosts, this is an issue and thus here we set out to use a single TetR-P_tet_^*^ system across various hosts. Nevertheless, the previous work with other versions of the promoter in various hosts highlights the potential of the system for use in even more bacterial phyla.

**Figure 2:**
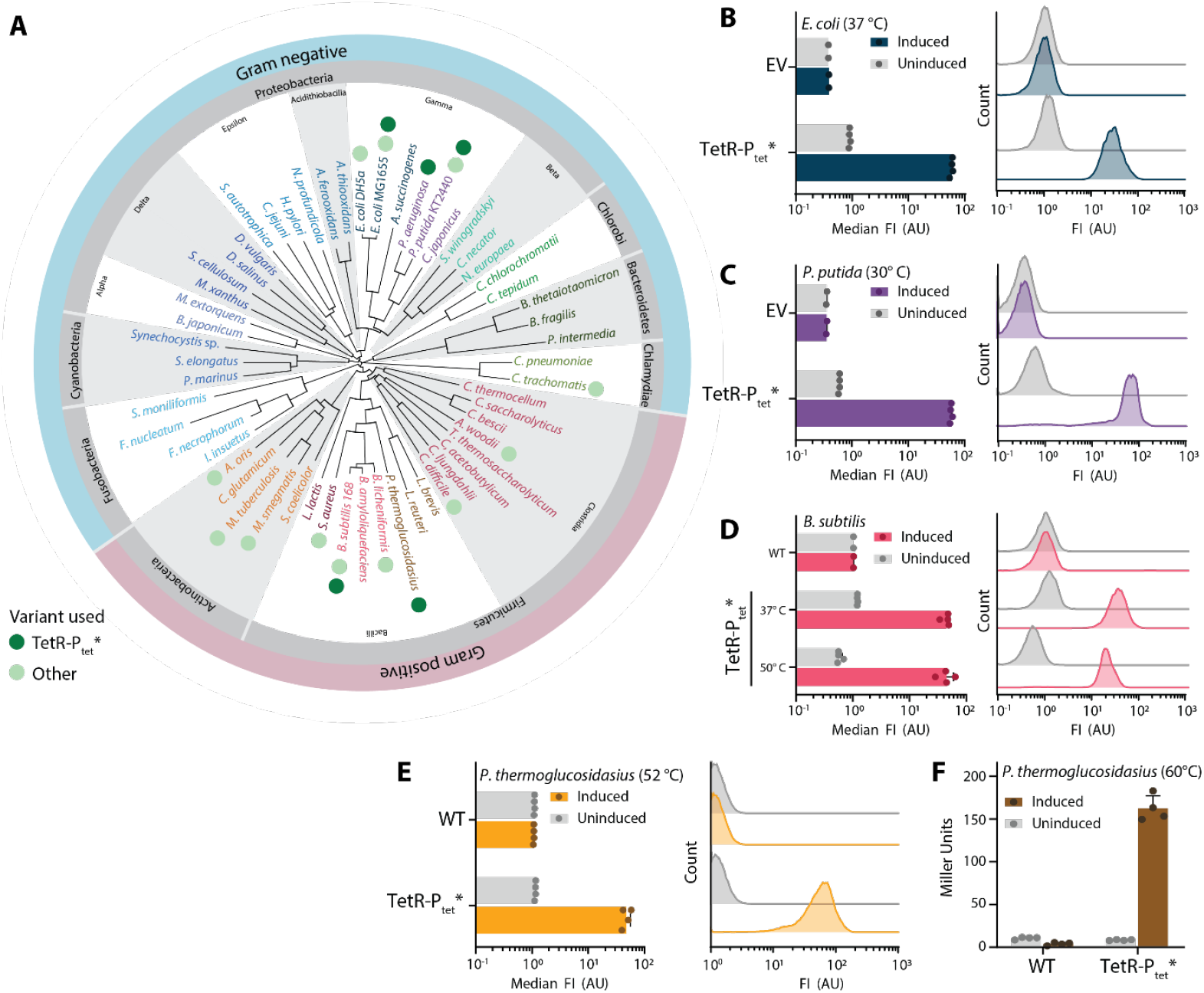
TetR-P_tet_^*^ allows broad-host-range application in industrially relevant prokaryotes. A phylogenetic tree summarizing organisms where, to our knowledge, various versions of the TetR-P_tet_ expression system are used (light green) and which organisms broad-host-range application of TetR-P_tet_^*^ is shown (dark green)(A). Single cell characterization of TetR-P_tet_^*^ controlling mRuby2 expression in *E. coli* (B), *P. putida* (C), *B. subtilis* (D) and *P. thermoglucosidasius* (E). *P. thermoglucosidasius* with TetR-P_tet_^*^ controlling *bgaB* expression at 60 °C is also shown (F). Either a strain containing an empty vector (EV) or wild type (WT) strain was used as control. Bar plots show the mean of median fluorescence intensities (FI) for biological quadruplicates or duplicates (for controls), with values of individual replicates indicated. For each condition, a histogram shows the single cell characterization for a representative replicate.

Though the inducer of the TetR-P_tet_ system, aTc, is generally less toxic than its antibiotic counterpart tetracycline, the structural similarities still result in toxicity in target organisms^52^. Therefore, initial toxicity screens of aTc on the chosen target organisms were performed. For each industrially relevant host, toxicity was observed at different levels, with *B. subtilis* being most sensitive to aTc concentrations of 0.1 µg mL^−1^ and *P. putida* displaying the highest tolerance to aTc, surpassing 2.0 µg mL^−1^ (Supplementary Figure S2).

Previously green, yellow and cyan fluorescent proteins have been engineered for improved thermostability^53^. However, considering the high autofluorescence of bacteria, particularly *Bacilli*, using a fluorescent protein in the red range (561 nm excitation) can allow higher resolution^22^. mRuby2 was previously shown to be a suitable fluorescent protein when a low signal-to-noise ratio of the system is crucial, and so was used for this study^22^. Using mRuby2 as reporter, at the same range of induction conditions, the titratability of the expression system across the hosts was determined. For *P. thermoglucosidasius* this was performed at 52 °C, as we observed that mRuby2 still functions at this temperature. To our knowledge, this is the first report using a red fluorescent protein in mild thermophilic conditions, which is useful to enable distinct fluorophores to be used simultaneously.

Investigating the titratability of the TetR-P_tet_^*^ system showed a linear range of induction within each host, though the dynamic range differs (Supplementary Figure S2). In *E. coli, B. subtilis* and *P. thermoglucosidasius*, aTc toxicity dictates maximum expression levels, whereas in *P. putida* promoter saturation at 1.0 µg mL^−1^ prevents a higher mRuby2 expression signal. With this analysis, optimum maximum induction conditions for *E. coli, P. putida, B. subtilis* and *P. thermoglucosidasius* were set at 1.0 µg mL^−1^, 1.0 µg mL^−1^, 0.1 µg mL^−1^ and 0.5 µg mL^−1^, respectively, and used for further experiments.

With suitable induction conditions identified, single cell characterization of the TetR-P_tet_^*^ system in the selected industrially relevant prokaryotes was performed. The results showed that the same TetR-P_tet_^*^ system is tight and homogeneous in all the tested hosts. Additionally, by performing time course experiments, we found that in all hosts a fast response time is observed, with maximum induction levels being reached within 2 to 4 hours of induction (the maturation time of mRuby2 is in the range of around 150 min^54^, Supplementary Figure S3). Additionally, the TetR-P_tet_^*^ system is uninduced and homogenous at mildly thermophilic temperatures (52 °C), as measured by the mRuby2 readout (Figure 2E), which is particularly interesting considering that the TetR-P_tet_^*^ is derived from a mesophilic host. To expand the use of the system to more thermophilic hosts with growth temperatures around 60 °C, a thermostable β-galactosidase (BgaB)^55^ was used as reporter, as mRuby2 fluorescence is no longer detectable at this temperature. At 60 °C, TetR-P_tet_^*^ response was still silent and inducible with a similar response time as seen at 52 °C (Figure 2F and Supplementary Figure S4). Consistent performance of the promoter in both mesophiles and thermophiles is noteworthy, as thermophilic expression cassettes often show inverted function, with either poor functionality in mesophiles and strong functioning in thermophiles or vice versa. This can particularly be an issue when cloning expression vectors, as high unintended expression in mesophilic cloning strains can place a significant mutational burden on the strain, and hence result in issues obtaining the desired vector and ultimately the desired production strain. Additionally, for thermophiles, there are not many inducible promoters available in the current toolbox, with the predominant reliance on (variants of) the xylose promoter^56,57^. Especially in the catabolism of alternative carbon sources, such as biomass hydrolysates, this may not be suitable. Therefore, the proposed TetR-P_tet_^*^ expression system is not only valuable for broad-host-range screening, but also for the use in thermophiles in general.

### Providing TetR-P_tet_^*^ in the proUSER2.0 vector collection allows interoperability

To facilitate the expansion of the range of organisms in which the TetR-P_tet_^*^ can be used, we provide a vector collection of broad-host-range plasmids harboring the expression system (Table 2). Within synthetic biology, various vector standards have been created that allow replication in specific subsets of prokaryotic hosts. Whereas the Standard European Vector Architecture^58^ (pSEVA) is the benchmark for Gram-negative hosts, the requirement of shuttle vectors limits its use in Gram-positive hosts. To overcome this, various shuttle vector standards that replicate in Gram-positive organisms have been developed, where the pMTL80000 system is most known for its replication across hosts^59^. Recently, a shuttle vector standard optimized for replication in *B. subtilis* was published^22^. Some of the vectors share dedicated origins of replication that have, with time, been shown to function in a wide range of organisms. The pBBR1 Gram-negative origin of replication was isolated from the *Bordetella bronchiseptica*^13^, and has since been widely applied to various Gram-negative hosts, particularly Proteobacteria. Though originating from the Gram-negative *Enterococcus faecalis*, the pAMβ1 origin of replication has been shown to function across Gram-positive hosts^60,61^. The pNW33n origin of replication (RepB) from *B. subtilis*^62^, has also been applied in synthetic biology, predominantly for use in *Bacilli*. With time, these origins of replications have been applied to a wide range of hosts, of which a comprehensive summary is shown in Table 3. However, though functional, it is important to note that the copy number of the plasmids harboring a particular origin of replication may vary between hosts^32^.

**Table 2:** Overview of the new ProUSER2.0 plasmids carrying the TetR-P_te_^*^ deposited at Addgene. Cam = chloramphenicol, Ery = erythromycin, Kan = kanamycin. Superscript “BS” designates that the integration locus is for *B. subtilis*. Superscript “Th” designates the thermostable variant of the kanamycin resistance cassette used in the rest of the collection, characterized by Y83D and K133T mutations^65^. The use of the Kan^Th^ variant or the TetR-P ^*^ variant is indicated with an “*”. When used as Gram-negative resistance marker lower concentrations of 6.25 μg mL^−1^ must be used. Supplementary Figure 5, provides a visual key, expanding on the previous ProUSER2.0 nomenclature^22^.

|  | Gram<sup>-</sup> Ab<sup>R</sup> | Gram<sup>-</sup> OriV | Gram<sup>+</sup> OriV or integration site | Gram<sup>+</sup> Ab<sup>R</sup> | Regulator & promoter |
| --- | --- | --- | --- | --- | --- |
| pProUSER63H0G | Gm<sup>R</sup> | pBBR1 | repB | Kan<sup>Th</sup> | TetR-P<sub>tet</sub><sup>*</sup> |
| pProUSER63I2G | Gm<sup>R</sup> | pBBR1 | pAMβ1 | Ery<sup>R</sup> | TetR-P<sub>tet</sub><sup>*</sup> |
| pProUSER63C1G | Gm<sup>R</sup> | pBBR1 | *gImS*<sup>BS</sup> | Cam<sup>R</sup> | TetR-P<sub>tet</sub><sup>*</sup> |
| pProUSER94H0G | Kan<sup>Th</sup> | ColE1 | repB | Kan<sup>Th</sup> | TetR-P<sub>tet</sub><sup>*</sup> |
| pProUSER94J0G | Kan<sup>Th</sup> | ColE1 | repB & *Δldh* | Kan<sup>Th</sup> | TetR-P<sub>tet</sub><sup>*</sup> |

**Table 3:** Overview of organisms with proven use of the broad-host-range origins of replications for Gram-negative hosts (BBR1) and Gram-positive hosts (pAMβ1 and the origin from pNW33N). Note the list is not exclusive, containing one example per organism, where more references may exist.

| Origin of replication | Phylum | Species |
| --- | --- | --- |
| *Gram-negative* |  |  |
| BBR1 | Actinomycetota | *Microbacterium* sp.<sup>67</sup> |
|  | Proteobacteria (Alpha) | *Acetobacter xylinum*<sup>14</sup>, *Agrobacterium tumefaciens*<sup>68</sup>, *Bartonella bacilliformis*<sup>14</sup>, *Brucella* spp.<sup>14</sup>, *Caulobacter crescentus*<sup>14</sup>, *Magnetospirillum gryphiswaldense*<sup>69</sup>, *Paracoccus denitrificans*<sup>14</sup>, *Rhizobium leguminosarum* by. *viciae*<sup>14</sup>, *Rhizobium meliloti*<sup>13</sup>, *Rhodobacter sphaeroides*<sup>14</sup>, *Sphingomonas* sp. LB126<sup>70</sup> |
|  | Proteobacteria (Beta) | *Alcaligenes eutrophus*<sup>14</sup>, *Bordetella pertussis*<sup>71</sup>, *Bordetella* spp<sup>13</sup>, *Burkholderia cenocepacia*<sup>72</sup>, *Burkholderia cepacia*<sup>73</sup>, *Cupriavidus metallidurans*<sup>74</sup>, *Ralstonia eutropha* H16<sup>75</sup> |
|  | Proteobacteria (Delta) | *Geobacter sulfurreducens*<sup>76</sup> |
|  | Proteobacteria (Gamma) | *Escherichia coli*<sup>13</sup>, *Pseudomonas fluorescens*<sup>14</sup>, *Pseudomonas putida*<sup>13</sup>, *Salmonella typhimurium*<sup>14</sup>, *Vibrio cholerae*<sup>13</sup>, *Xanthomonas campestris*<sup>14</sup>, *Salmonella enterica*<sup>77</sup>, *Pseudomonas aeruginosa*<sup>78</sup>, *Edwardsiella ictaluri*<sup>79</sup> |
| *Gram-positive* |  |  |
| pAMβ1 | Actinomycetes | *Brevibacterium lactofermentum*<sup>80</sup>, *Brevibacterium flavum*<sup>80</sup>, *Corynebacterium glutamicum*<sup>80</sup>, *Corynebacterium melassecola*<sup>80</sup> |
|  | Firmicutes (Bacilli) | *Bacillus subtilis*<sup>60</sup>, *Bacillus thuringiensis*<sup>81</sup>, *Enterococcus faecalis*<sup>61,82</sup>, *Lactobacillus acidophilus*<sup>83</sup>, *Lactobacillus reuteri*<sup>83</sup>, *Lactobacillus salivarius*<sup>83</sup>, *Lactococcus lactis*<sup>81</sup>, *Listeria monocytogenes*<sup>81</sup>, *Staphylococcus aureus*<sup>81</sup>, *Streptococcus agalactiae*<sup>81</sup> |
|  | Firmicutes (Clostridia) | *Clostridium acetobutylicum*<sup>84</sup>, *Clostridium beijerinckii*<sup>85</sup>, *Clostridium cellulolyticum*<sup>86</sup> |
| pNW33N_ori (RepB) | Firmicutes (Bacilli) | *Bacillus coagulans*<sup>87</sup>, *Bacillus methanolicus*<sup>53</sup>, *Bacillus smithii*<sup>53</sup>, *Bacillus subtilis*<sup>88</sup>, *Bacillus thuringiensis*<sup>89</sup>, *Geobacillus stearothermophilus*<sup>90</sup>, *Geobacillus thermodenitrificans*<sup>53</sup>, *Paenibacillus polymyxa*<sup>91</sup>, *Parageobacillus thermoglucosidasius*<sup>92</sup>, *Staphylococcus aureus*<sup>93</sup> |
|  | Firmicutes (Clostridia) | *Clostridium thermocellum*<sup>94</sup> |

Here, we built the vector collection on the premise created by the ProUSER2.0 system^22^, though the origins of replication and selection cassettes of choice allow its use in other Gram-positives, as indicated in Table 3. Additionally, as the ProUSER system has previously been used for Gram-negatives, it is a suitable platform to build upon^63^. For the characterization done here, one broad-host-range SD sequence and spacer were selected, however, with the ProUSER2.0 cloning system, easy exchange of SD sequence and spacer length can be achieved when this needs to be tailored for specific purposes.

In the collection, pProUSER63H0G is provided that allows broad-host-range expression from a single vector in a wide range of mesophiles and thermophiles. As there is a limited number of resistance markers that can be used in thermophiles, we provide pProUSER63I2G to allow broad-host-range expression amongst mesophiles. When *B. subtilis* is selected as desired production host, pProUSER63C1G can be used to simply create more stable strains through integration into the genome at the previously reported, favorable *glmS* integration site^22^. Considering that the TetR-P_tet_^*^ system is in itself an advantageous expression system in thermophiles, we further provide pProUSER94H0G and pProUSER94J0G. pProUSER94H0G can be used for replicative screening of constructs in thermophiles and pProUSER94J0G can be used to integrate the constructs replacing the lactate dehydrogenase (LDH) gene of the model thermophile *P. thermoglucosidasius* to create more stable expression strains. This locus was selected since lactate is the main byproduct formed in this strain, hampering efficient pathway yields of other target products and therefore typically an advantageous gene to delete in metabolic engineering projects^64^. With the premise of the ProUSER2.0 system, and the application module being transferable to both the pSEVA collection and the pMTL80000 systems, modifications to these vectors can easily be made to tailor them to a specific host, once screening has identified its endogenous metabolism as most favorable.

### TetR-P_tet_^*^ is a valuable tool in bacterial models of pathogens

Aside from being useful as a broad-host-range promoter for industrially relevant prokaryotes, we expanded the host range application of the TetR-P_tet_^*^ expression system for other purposes. As *Pseudomonas aeruginosa* is an often used model host for chronic lung infections, demonstrating functioning in this host can expand the use to more mechanistic medically relevant approaches as well. To test the functionality of the TetR-P_tet_^*^ expression system, the same plasmid as tested in the industrially relevant hosts (pVM62), was used to evaluate the performance through mRuby2 readout (Figure 3). Similar to the industrially relevant hosts, the TetR-P_tet_^*^ system was silent, titratable and homogeneous across the population in *P. aeruginosa*. The lowered sensitivity to aTc as an inducer shows the importance of fine tuning the inducer concentration when the expression system is tested in a new host. *P. aeruginosa* has been shown to possess a large array of efflux pumps^66^, resulting in lowered sensitivity to the aTc inducer. Therefore higher concentrations of aTc are needed before the activation and titratable change of the expression as measured through mRuby2 output is observed (Figure 3A, B). However, when adequate aTc concentrations are used, the titratable, homogeneous and timely response of the expression cassette can still be observed (Figure 3C, Supplementary Figure S6). We provide various plasmids (pProUSER63H0G-mRuby2, pProUSER94H0G-mRuby2 and pVM62) to allow simple validation and further use in alternative desired target hosts.

**Figure 3:**
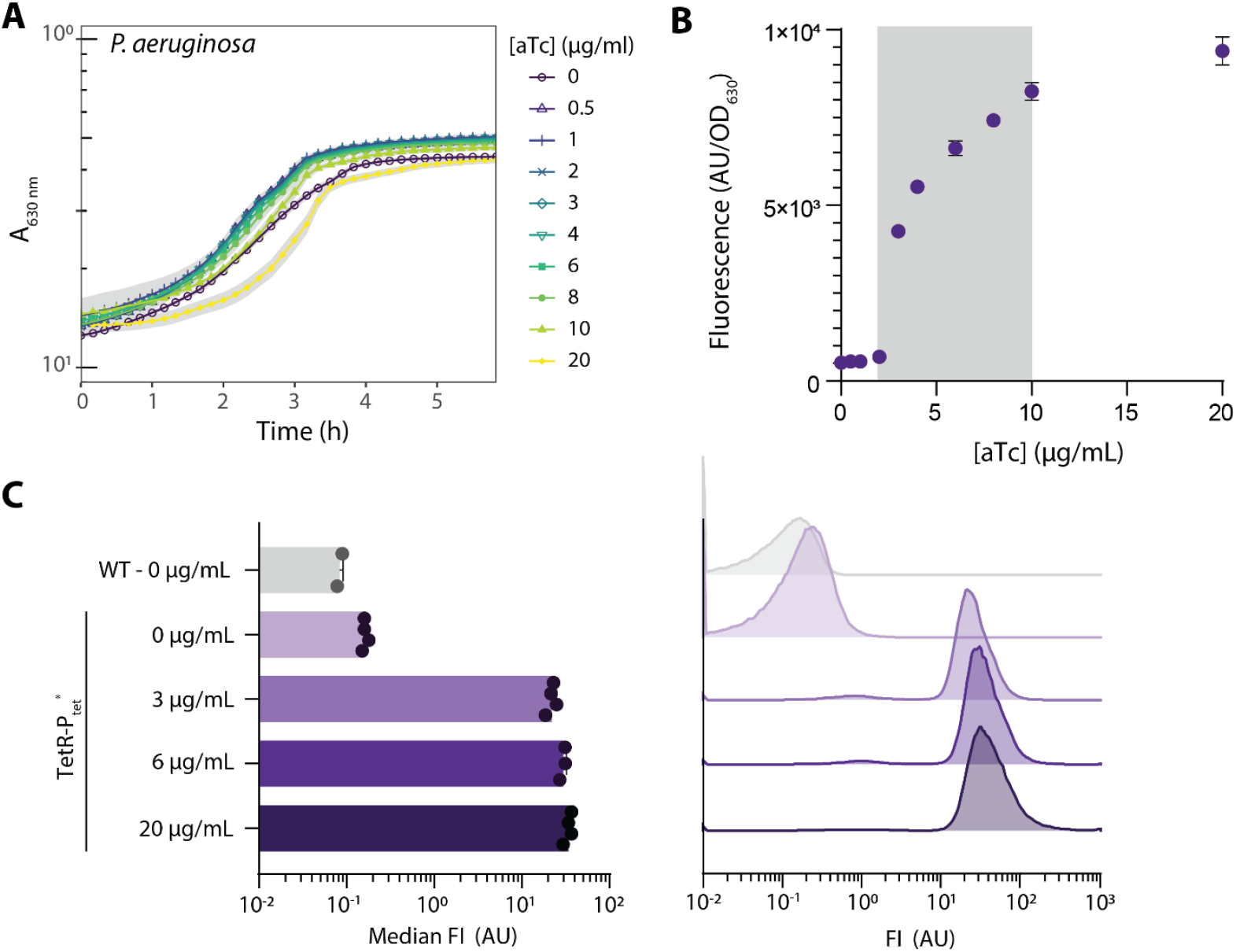
TetR-P_tet_^*^ functioning in *P. aeruginosa*, a model organism for pathogenic species. Microtiter plate reader measurements show the toxicity (A) and titratability (B) of the system for different aTc concentrations. Time course shake flask experiments show the homogeneity of the expression system for varying aTc concentrations used (C), with a representative histogram of each condition shown on the right. All data is shown in biological duplicates or quadruplicates, with error bars indicating the standard deviation of measurements.

### TetR-P_tet_^*^ allows screening of protein functionality across hosts

Phenolic compounds form a wide range of organic compounds, with varying applications in the pharmaceutical industry, food industry as well as in the production of bioplastics^95^. Examples of these compounds include but are not limited to, *p-*coumaric acid, caffeic acid and ferulic acid, all derivatives of the aromatic amino acid tyrosine. Enzymes that can interconvert these and other phenolic compounds have predominantly been found in plants, cyanobacteria and actinomycetes. Therefore, expression of these proteins in microbial prokaryotic hosts for bioproduction or protein assays can be challenging in the predominantly chosen *E. coli* host^96,97^. To increase the expression of these enzymes, allowing better bioproduction as well as enzymatic studies, finding more suitable microbial hosts, without the need to heavily engineer the enzyme, can be valuable. Here, we demonstrate how the TetR-P_tet_^*^ system can efficiently be used to screen three different enzymes, all producing a phenolic compound in various mesophilic hosts by only creating one vector per enzyme. The first enzyme selected is 4-Coumarate 3-Hydroxylase (C3H) from *Saccharothrix espanaensis*, catalyzing the flavin adenine dinucleotide (FAD) dependent conversion of *p*-coumaric acid into caffeic acid (Figure 4A), contrary to the C3H P450 enzymes that require heme as co-factor^98^. Caffeic acid can be methylated into ferulic acid by the action of caffeic acid O-methyltransferase (COMT) from *Arabidopsis thaliana*, which through codon optimization was previously shown to be expressed in engineered *E. coli*^99^. Previously an O-methyltransferase from the cyanobacterium *Synechocystis* sp. (SynOMT), was identified which allows a wider range of catalytic transformations. Both COMT and SynOMT are dependent on the common methyl group donor S-adenosyl-L-methionine (SAM). Considering that cofactor availability and regulation vary greatly between organisms, protein expression can benefit from screening across microbial hosts.

**Figure 4:**
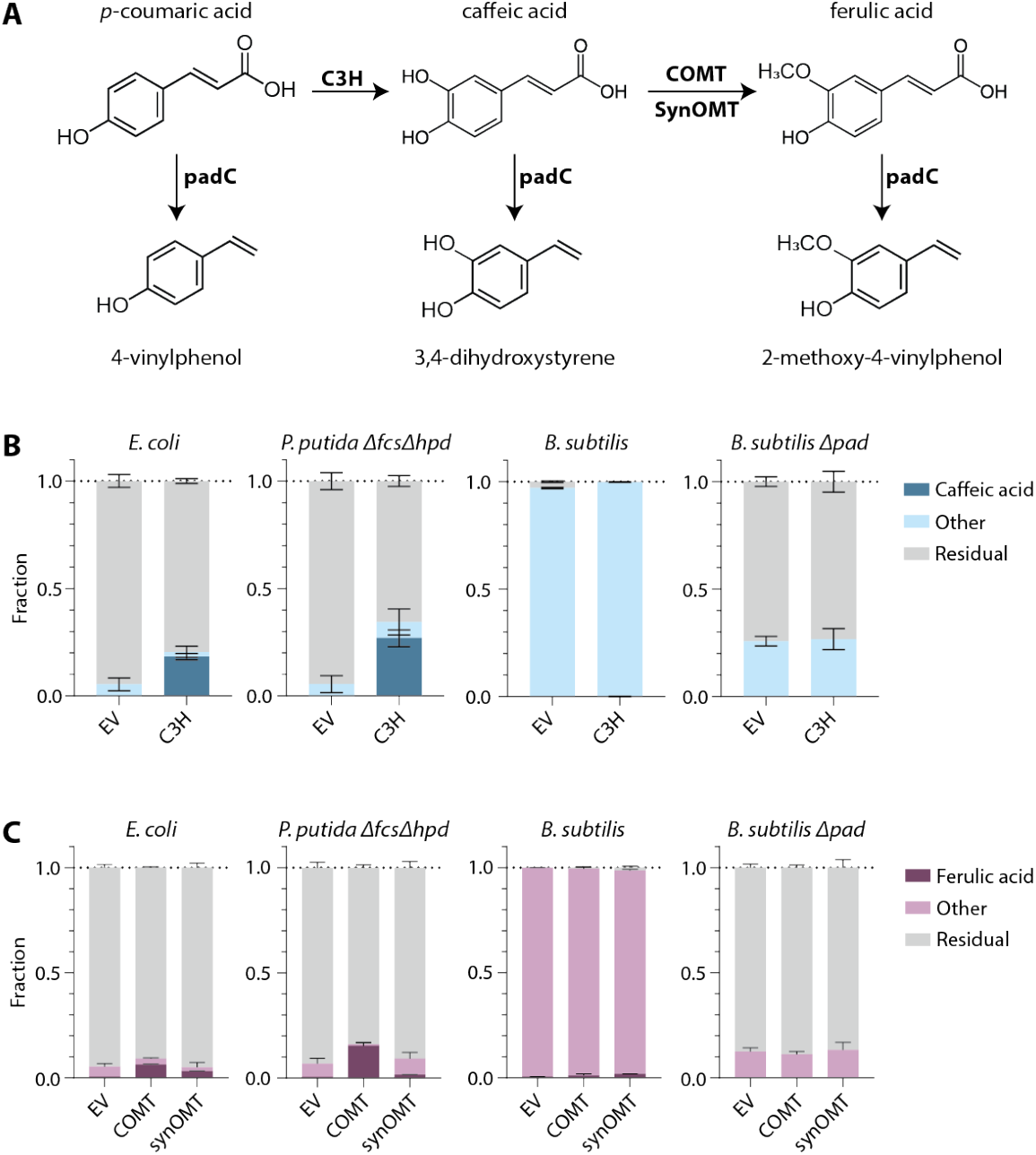
Screening of mesophilic hosts for analyzing protein functionality of a 4-Coumarate 3-Hydroxylase (C3H) and two caffeic acid O-methyltransferases (COMT and synOMT). Overview of the reactions catalyzed by the enzymes of interest, as well as a phenolic acid decarboxylase (PadC) (A). The fraction of the fed precursor converted 22 hours post induction is shown when *p*-coumaric (B) or caffeic acid (C) are fed to *E. coli, P. putida* Δ*fcs* Δ*hpd, B. subtilis* and *B. subtilis* Δ*pad* cultures. All data is shown in biological quadruplicates, with error bars indicating the standard deviation of measurements.

To screen the expression of C3H, COMT and synOMT in various hosts, we inserted the coding sequences of the genes into pProUSER63H0G and used one plasmid per enzyme to screen expression in *E. coli, P. putida* and *B. subtilis*. Considering that all enzymes are of mesophilic origin, and no data is available on the potential thermostability of these enzymes, they were only screened in mesophilic hosts. *P. putida* has previously been reported to be able to degrade *p*-coumaric acid, and so a previously engineered *P. putida* KT2440Δ*fcs*Δ*hpd*, lacking this ability, was used as background^63^. Additionally, the presence of a phenolic acid decarboxylase (PadC) in *B. subtilis* results in the conversion of a wide range of phenolics into unwanted side products. PadC can for example decarboxylate *p*-coumaric acid, caffeic acid, and ferulic acid into 4-vinylphenol, 3,4-dihydroxystyrene, and 2-methoxy-4-vinylphenol, respectively (Figure 4A)^100^. As this could affect conversion by the expressed C3H, COMT and SynOMT enzymes, we engineered a *B. subtilis* Δ*pad* strain. To analyze the efficiency of enzyme expression, conversion of the precursor compound was measured as the output of catalytic efficiency, the predominantly important factor in strain design. The concentration of precursor added was determined by toxicity studies across the various hosts (Supplementary Figures S7 and S8). From the screening of a single plasmid across the hosts, we found that *P. putida* Δ*fcs*Δ*hpd* was the most effective host for catalytic conversion by C3H and COMT, whereas *E. coli* showed the best conversion when expressing the synOMT enzyme, although overall conversion rates remained low (Figure 4B, C). Considering the similar functionality of the OMT enzymes, these results show the importance of screening enzymes across microbial hosts. Due to the presence of PadC, *B. subtilis* converts all of the fed *p*-coumaric or caffeic acid into 4-vinylphenol or an unknown compound, respectively. Through analysis of 3,4-dihydroxysytrene and 2-methoxy-4-vinylphenol standards on the HPLC, it was concluded that these compounds were not accumulating when *B. subtilis* was fed with caffeic acid. Accumulation of these compounds are also not observed when the *pad* knock out was introduced, however production of caffeic or ferulic acid was also not detected. These experiments highlight the applicability and ease of using the TetR-P_tet_^*^ system to create a single plasmid that allows screening of protein functionality across a series of prokaryotic hosts.

## Discussion

Through the development of the synthetic biology field, an increasing number of tools that enable the engineering of novel organisms have emerged. However, empirical preference for a small subset of model organisms still dictates host selection for many applications. To facilitate a more systematic exploration of the microbial phenotype landscape without requiring *a priori* knowledge of metabolism, previous work on broad-host-range tools has focused on the development of backbone vectors, specifically the origin of replications and selection cassettes. To further push the field towards allowing endogenous metabolism to guide host selection, we here present a broad-host-range inducible expression system, based on the well-known TetR-P_tet_ system though with significant modification to permit broad-host-range functionality. Through modification of the TetR-P_tet_ promoter, we show that it is tight, titratable and homogeneous across various prokaryotic hosts with industrial potential as well as model organisms with medical relevance. Additionally, the system is thermostable up to 60 °C, providing a novel tool for more controlled expression in thermophilic hosts. To expand the use of the TetR-P_tet_^*^ system past the organisms present here, we provide pVM62, which can easily be used to evaluate the system functioning through mRuby2 expression. We further demonstrate how endogenous physiology can influence functionality through the expression of a hydroxylase and two methyltransferases, allowing the selection of a more suited industrially relevant host as well as easier screening across multiple hosts for patenting purposes.

Though the TetR-P_tet_^*^ system proposed here has desirable characteristics in a wide range of hosts, when comparing protein expression levels, unintentional and uncontrolled bias towards host specific expression machinery may be present. For the system used here, for example, an SD sequence and spacer region have been selected with *a priori* knowledge from studies done in specific target hosts. Though the selected sequence allows functioning across species, tweaking this further could introduce slight expression bias in certain hosts. Additionally, the codon usage of the desired construct can influence protein expression and folding between hosts. However, since the predictive capabilities of these factors are limited, they are difficult to adjust for in current design workflows. To our knowledge, to date, there are no studies that investigate the design space of these components in more than two phylogenetically distinct prokaryotes. Transferring this context dependent knowledge into a predictive tool could allow further optimization of the TetR-P_tet_^*^ sequence to remove some of the unintentional bias that is currently expected to be present. Though an unintentional host bias may be present, the TetR-P_tet_^*^ system provided here forms a suitable platform to perform more context dependent comparative studies across organisms to help elucidate and understand the differences between functional parts. Because of the complexity of the parts, having the single TetR-P_tet_^*^ system functioning across hosts is a valuable tool for initial studies of expression and production across hosts.

Using the TetR-P_tet_^*^ system can enable the selection of a more suitable starting host for expression, functionality or production studies. However, upon initial screening, further tailoring of the system is necessary to achieve desired final characteristics. For example, when producing a desired protein of interest, integrative expression can be critical to ensure additional stability and homogeneity in a population. Further fine-tuning of the coding sequence or control elements may also be needed. Providing the TetR-P_tet_^*^ system embedded in the proUSER2.0 system offers easy exchange of SD and spacer sequences, as well as fine-tuning the target construct, as was previously demonstrated when screening a signal peptide library with the system^22^. For large scale production, the addition of an (expensive) inducer compound is typically avoided, and the selection of an auto-inducible or constitutive expression system can be more beneficial when working at larger scale. The power of the TetR-P_tet_^*^ system thus predominantly lies in initial screening and comparative studies of target hosts, albeit for a wide range of purposes.

When screening various hosts, transformation protocols for all desired organisms are undoubtedly required. Fortunately, with the expansion of synthetic biology tools, transformation protocols for many hosts have been reported in various publications^57,101–103^. Additionally, detailed and descriptive protocols have been exceedingly deposited in repositories, such as OpenWetWare^104^, making the transformation of a wide range of prokaryotes accessible. Increased focus on data and protocol availability in the field will only enhance this moving forward. Therefore, the TetR-P_tet_^*^ platform provided here can enable a valuable shift in synthetic biology steering away from single host based studies and applications, to a broader utilization of the microbial physiologies provided by nature through evolution, to society’s full advantage.

## Methods

### Strains, media, and growth conditions

The strains used in this study are summarized in Supplementary Table S1. *E. coli* K-12 MG1655, *B. subtilis* 168 and *P. aeruginosa* PAO1 strains were routinely grown in LB media at 37 °C with shaking at 250 revolutions per minute (RPM), or on agar plates at 37 °C according to standard protocols^105^. *P. putida* KT2440 was routinely grown on the same media, but at 30 °C with 250 rpm. *P. thermoglucosidasius* DSM2542^T^ was routinely grown on 2-SPY medium^106^ at 60 °C with 150 RPM shaking or on Tryptic Soy Agar plates (BD Biosciences) at 60 °C. *E. coli c*ultures used for plasmid prep were grown in media with double the amount of tryptone and yeast extract. When appropriate, media was supplemented with antibiotics (10 μg mL^−1^ gentamycin (Gm) or 6.25 μg mL^−1^ kanamycin (Km) for *E. coli* strains, 10 μg mL^−1^ Gm for *P. putida* and *P. aeruginosa*, 10 μg mL^−1^ erythromycin (Ery) for *B. subtilis* and 12.5 μg mL^−1^ Km for *P. thermoglucosidasius* strains).

For *E. coli, P. putida*, and *B. subtilis* characterization experiments were performed in modified minimal M9 medium, named M9extra (12.8 g L^−1^ Na_2_HPO_4_·7H_2_O, 3 g L^−1^ KH_2_PO_4_, 0.5 g L^−1^ NaCl, 1 g L^−1^ NH_4_Cl, 2 mM MgSO_4_, 0.1 mM CaCl_2_) supplemented with 60 μM FeCl_3_, a trace element solution (1.25 μM MnCl_2_·4H_2_O, 0.21 μM CoCl_2_·6H_2_O, 0.85 μM ZnSO_4_·7H_2_O, 0.05 μM CuCl_2_·2H_2_O, 0,08 μM H_3_Bo_3_, 0.105 μM NiCl_2_·6H_2_O, 0.125 μM NaMoO_4_·2H_2_O) and 10 g L^−1^ D-glucose as carbon source, whereas LB was used for *P. aeruginosa*. For *P. thermoglucosidasius* characterizations, Thermophile Minimal Medium (TMM) was used, adapted from Fong et al.,(2006), and contains, per liter: 930 ml Six salts solution (SSS), 40 ml 1 M MOPS (pH 8.2), 10 ml 1 mM FeSO_4_ in 0.4 M tricine, 10 ml 0.132 M K_2_HPO_4_, 10 ml 0.953 M NH_4_Cl, 0.5 ml 1 M CaCl_2_, 1x Wolfe’s vitamin solution and trace element solution, adjusted to a final pH of 6.8. SSS contained, per 930 ml: 4.6 g NaCl, 1.35 g Na_2_SO_4_, 0.23 g KCl, 0.037 g KBr, 1.72 g MgCl_2_·6 H_2_O and 0.83 g NaNO_3_. The trace element solution contained, per liter, 1 g FeCl_3_·6 H_2_O, 0.18 g ZnSO_4_·7 H_2_O, 0.12 g CuCl_2_ ·2 H_2_O, 0.12 g MnSO_4_·H_2_O and 0.18 g CoCl_2_·6 H_2_O. 1000x Wolfe’s vitamins consist of, per liter, 10 mg pyridoxine hydrochloride, 5.0 mg thiamine-HCl, 5.0 mg riboflavin, 5.0 mg nicotinic acid, 5.0 mg calcium D-(+)-pantothenate, 5.0 mg p-aminobenzoic acid, 5.0 mg thioctic acid, 2.0 mg biotin, 2.0 mg folic acid and 0.1 mg vitamin B12. A Final of 0.2% yeast extract and 10 g L^−1^ D-glucose were supplemented to the medium.

For *E. coli, P. putida* and *B. subtilis* production experiments, M9extra medium with 20 g L^−1^ D-glucose and 0.1% yeast extract (YE) was used. Additionally, for *B. subtilis* experiments a final concentration of 50 mg L^−1^ L-tryptophan was also added.

### PCR amplification, DNA purification, and sequencing

PCR amplification for USER cloning was performed using Phusion U (Thermo Fisher Scientific), where Phusion HSII (Thermo Fisher Scientific) was used for sequencing of genomic DNA integrations and OneTaq was used for colony PCRs. *E. coli, P. putida* and *P. aeruginosa* colony PCRs were performed by suspending a colony in 20 µL LB media and using 1 µL of this suspension as the template in the reaction mixture. For colony PCR of *B. subtilis* strains, individual colonies were suspended in 20 µL LB media, and 10 µL of this suspension was diluted in 10 µL MiliQ water. The dilutions were stored on ice for 5 minutes, followed by 3 cycles of 1 minute microwave heating at 800W and 30 seconds on ice. After cooling the dilutions on ice for 5 minutes, 1 µL of the dilution was used as the template in the reaction mixtures. For *P. thermoglucosidasius* colony PCR, individual colonies were suspended in 20 µL 20mM NaOH and subjected to subsequent boiling at 95 °C for 15 minutes. After cooling to room temperature, 1 µL of the mixture was used as the template in the reaction mixture.

The modified TetR-P_tet_ regulator and promoter sequence was ordered from IDT Technologies as Gblock, and designed using the pProUSER13G2F sequence as template^22^, with specific point mutations introduced to remove the identified and undesired restriction sites.

A NucleoSpin PCR clean-up gel extraction kit (Macherey-Nagel, Germany) was used to purify PCR products before their use in cloning, according to the manufacturer’s instructions. Plasmid DNA was purified from *E. coli* cultures using a NucleoSpin Plasmid kit (Macherey-Nagel, Germany), following the manufacturer’s instructions. Routine sequencing was performed using a Mix2Seq kit (Eurofins Genomics, Luxembourg), following the manufacturer’s instructions.

### Plasmid construction

The vectors used in this study for characterization of the TetR-P_tet_ promoters were constructed using USER cloning. For the USER reactions, the different fragments were mixed with 1.2 µL T4 ligase buffer (New England BioLabs, United States), 1 μL USER enzyme (New England BioLabs, United States), and DNAse/RNAse free miliQ to a total volume of 12 μL. The USER reactions were incubated at 37 °C for 25 minutes, followed by 25 minutes at 25 °C. Subsequently, the reactions were diluted by the addition of 8 μL DNAse/RNAse free MilliQ, and 5 μL was used for transformation.

USER ligation PCR was used to generate a linear fragment to knock out the phenolic acid decarboxylase (*pad*, NCBI Ascension number AF017117) from *B. subtilis*. After the USER reaction with desired fragments, 6.2 µL of MilliQ, 0.8 µL T4 DNA ligase buffer and 1 µL T4 DNA ligase (Sigma) were added. After incubation for at least 1 hour at room temperature, the ligated product was amplified by PCR by using 1-2 µL of the reaction mixture.

To insert the target proteins into the created proUSER2.0 vectors, backbone digestion was performed according to previously described protocols^22^. The insert genes were PCR amplified using primers to generate an AGGAGG ribosome binding site and a 7 bp spacer. Transformation of the USER reactions into *E. coli* DH5α-λ***pir*** was performed according to standard protocols^107^. The list of constructed plasmids and primers used in this study can be found in Supplementary Tables S2 and S3 respectively. Plasmid maps of the used plasmids are attached as Supplementary Data.

### Transformations of bacterial cells & strain construction

Plasmids were transformed into *E. coli, P. putida* and *P. aeruginosa* through standard electroporation protocols^103^. To transform *B. subtilis* glms::P_mtlA_-comKS, the engineered competence was induced according to previously published protocols^108^. *P. thermoglucosidasius* transformation was performed as described previously^57^. To integrate constructs into the *P. thermoglucosidasius* genome, a transformant was inoculated into 2-SPY with 12.5 μg mL^−1^ kanamycin and passaged serially at 60 °C with 150 RPM for 1-2 days after which a dilution plated on TSA with 12.5 μg mL^−1^ kanamycin. Resistant colonies were screened for proper first integration into the desired locus, and subsequently inoculated into 2-SPY medium for serial passaging at 65 °C, 150 RPM for 2-5 days. Replica plating on TSA and TSA with 12.5 μg mL^−1^ kanamycin revealed sensitive colonies, of which the stable integration was confirmed with colony PCR and sequencing.

### Toxicity and inducibility characterizations

To determine the toxicity of aTc and the inducibility of the TetR-P_tet_ promoter in the various organisms, a colony was inoculated into 5 mL rich medium, and grown for 8 hours at appropriate temperature and agitation. 50 µL of this rich pre-culture was used to inoculate 5 mL minimal medium for overnight growth. The overnight pre-culture was used to inoculate a clear-flat-bottomed 96-well microtiter plate with the minimal medium at a starting Optical Density (OD_600_) of 0.1, at a volume of 190 µL. After 2 hours from inoculation, cultures were induced with varying levels of aTc, as indicated, and growth and fluorescence were monitored with a Synergy HM1 absorbance and fluorescence reader (BioTek Instruments, United States). For *E. coli, P. putida* and *P. aeruginosa* strains, the plate was grown in the plate reader, and OD_600_ and mRuby2 fluorescence (λ_ex_ = 559 nm, λ_em_= 600nm) was measured every 15 minutes for a minimum of 14 hours. For *B. subtilis* and *P. thermoglucosidasius* the inoculated plate was placed in a 37 °C or 52 °C incubator, respectively, and manually measured every 2 hours for 12 hours.

To determine the toxicity of the precursor chemicals, caffeic and *p*-coumaric acid, 10 µL of an 8h culture in LB medium was used to seed 50 mL M9extra medium with 10 g L^−1^ D-glucose and 0.05% YE, with an additional 50 mg mL^−1^ L-trytophan for *B. subtilis*. 5 μL of the overnight culture was transferred to 95 μL M9extra media with 5 g L^−1^ D-glucose, 0.05% YE (+ 50 mg·L-1 L-tryptophan for *B. subtilis*) in a 96-well microtiter plate and incubated for 2 hours at either 37 °C or 30 °C in a plate reader with shaking. 100 μL of pre-heated serial dilutions of *p*-coumaric acid and caffeic acid dissolved in culture media was added and growth was followed for 24 hours by 15 minute interval measurements at 630 nm with an ELx808™ Absorbance Microplate Reader.

### Flow cytometry characterization

For single-cell characterization of the TetR-P_tet_ promoters, a single colony of the strains were inoculated into 1 mL rich media (LB or TSA), and grown for 8h at the optimum temperature and appropriate agitation. 10 µL of the preculture was used to inoculate 50 mL of minimal medium, supplemented with 0.2% YE, and grown overnight. The final characterization cultures were inoculated at a starting OD_600_ of 0.05 and induced with the suitable level of aTc after the culture had grown to an OD_600_ of at least 0.1. OD_600_ was monitored with a Jenway 6705 spectrophotometer. 1 mL samples for flow cytometry analyses were taken and centrifuged at 6000g for 10 minutes. The supernatant was exchanged with 500 µL 2% paraformaldehyde in phosphate-buffered saline (PBS) (Thermo Fisher Scientific, United States). The samples were left at room temperature for 30 minutes to 2 hours for fixating, and the fixated cells were subsequently stored in PBS at 4 °C until analysis within 2 weeks of sampling.

The fluorescence of the fixed cells was quatified using a MACSQuant VYB cytometer (Miltenyi Biotec, Germany). mRuby2 fluorescence was measured by excitation with a 561 nm laser, using a 615/20 nm band-pass filter. Per 200 µL sample, up to 50,000 events were recorded using varying gain settings per organism measured. For *E. coli* and *P. putida*: forward scatter: 718V, side scatter: 570 V, Y2 channel: 450 V. For *B. subtilis* and *P. thermoglucosidasius*: forward scatter: 580 V, side scatter: 516 V, Y2 channel: 492 V. For *P. aeruginosa*: forward scatter: 500 V, side scatter: 450 V, Y2 channel: 399 V. Data was analyzed using Flowjo (Tree Star Inc., Asland, OR).

### B-galactosidase assay

To characterize the TetR-P_tet_^*^ promoter in *P. thermoglucosidasius* at 60 °C, a thermostable β-galactosidase^55^ was used as a replacement for the mRuby2 reporter. Growth experiments were performed in the same manner as the flow cytometry characterization. Upon sampling, a Miller assay was performed to quantify β-galactosidase activity, according to a previously described protocol^109^, with one modification. The enzymatic reactions proceeded at 65 °C, the optimum temperature observed for the thermostable β-galactosidase. One Miller unit represents a standardized amount of β-galactosidase. The total Miller units per sample were found by measuring the absorbance at 420 nm (A_420nm_), the reaction time (t) and total assay volume (v), the sample OD (A_600nm_) and calculated by:

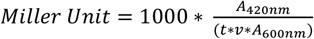

### Protein functionality assessment

To determine the *in vivo* functionality of the 4-coumarate 3-hydroxylase (C3H) and the caffeic acid O-methyltransferase ((syn)OMT), overnight cultures were diluted to an OD_600_ of 0.05, after which 2 mL was added to a 24-deep well plate. After 2 hours of incubation, cultures were induced by the addition of aTc at the indicated concentrations. *E. coli* and *P. putida* cultures were grown at 30 °C, whereas *B. subtilis* cultures were grown at 37 °C. 0.5 hours after induction, 2 mL of the culture medium supplemented with 5 mM *p*-coumaric acid or 20 mM caffeic acid and appropriate aTc concentration was added to each well. 700 µL samples were taken 22 hours post induction to measure culture OD, after which the sample was centrifuged at 6500g and sample supernatants were diluted 1:1 with absolute ethanol and used for HPLC analysis. *p*-Coumaric acid, caffeic acid, 4-vinylphenol, 3,4-dihydroxystyrene and 2-methoxy-4-vinylphenol concentrations were quantified using a Dionex UltiMate 3000 HPLC (Thermo Scientific) with a Discovery® HS-F5-3 column (15 cm x 2.1 mm, 3 μm particle size, Supleco Analytical). 2.5 µL sample was injected into the column, and kept at 30 °C. A constant flow rate of 0.7 mL min^−1^ was used with a linear gradient of two mobile phases, 10 mM ammonium format (pH adjusted to pH 3.0 with formic acid, phase A) and acetonitrile (phase B), as follows: 0 to 1.0 min (10% B), 1.0 to 7.5 min (linear gradient from 10 to 90% B), 7.5 to 9.0 min (90% B), 9.0 to 9.2 min (90 to 10% B) and 9.2 min to 12.0 min (10% B). The compounds were detected at a wavelength of 280 nm.

### Phylogenetic analysis

To generate the phylogenetic tree, 16S rRNA sequences from various hosts across bacterial phyla were obtained from the National Center for Biotechnology Information (NCBI). A list of all strains used with corresponding NCBI accession numbers is given in Supplementary Table S4. ClustalW with character counts was used and visualized with plotTree in R^110,111^.

## Author Contributions

VM and SIJ conceived the study and designed the experiments. VM, KBF, ADAW, IP, CB and ALS performed the experiments. VM and SIJ analyzed the data and wrote the manuscript. Supervision was done by ATN and SIJ. All authors have read, corrected and approved the manuscript.

## Acknowledgments

This study was funded by the Copenhagen Bioscience PhD Program (Grant no. NNF18CC0033664; VM) and the Independent Research Foundation Denmark (Grant no. 7017-00321B; SIJ). We further acknowledge funding from the Novo Nordisk Foundation (Grant no. NNF20CC0035580) and European Union’s Horizon 2020 research and innovation program under Grant Agreement No. 814408 (SHIKIFACTORY100).

## Conflicts of interest

The authors declare no competing financial interest.

## Notes

### Competing Interest Statement

The authors have declared no competing interest.

