## Supplementary Figures and Tables for "From empirical to data-driven host selection: a broad-host-range expression platform to facilitate chassis screening"

### List of content

Supplementary Figure S1: Comparison of mRuby2 expression controlled by either TetR-$P_{tetA}$ andTetR-$P_{tet*}$

Supplementary Figure S2: Toxicity of aTc and titratability of TetR-$P_{tet}$*

Supplementary Figure S3: Time course experiment using flow cytometry for quantification of mRuby2 expression under the control of tetR-$P_{tet}$* in different relevant cell factories

Supplementary Figure S4: Time course experiments of *P. thermoglucosidasius* expressing BgaB under the control of tetR-$P_{tet}$* at 60°C

Supplementary Figure S5: Time course experiment using flow cytometry for quantification of mRuby2 expression under the control of tetR-$P_{tet*}$ in *P. aeruginosa*

Supplementary Figure S6: Overview of the updated ProUSER2.0 nomenclature

Supplementary Figure S7: Toxicity of p-Coumaric acid in different potential cell factories

Supplementary Figure S8: Toxicity of caffeic acid in different potential cell factories

Supplementary Table S1: Strains used in this study

Supplementary Table S2: Plasmids used in this study

Supplementary Table S3: Oligos used in this study

Supplementary Table S4: 16S sequences from corresponding strains used for generating the phylogenetic tree

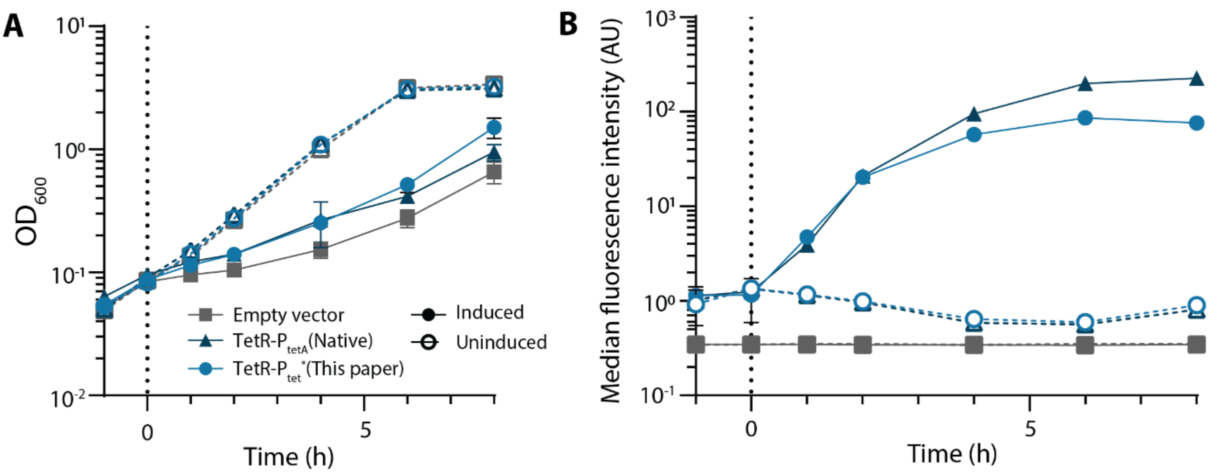


**Supplementary Figure S1:** Time response of *E. coli* cultures expressing mRuby2 under the control of either the TetR-P$_{tetA}$ or TetR-P$_{tet*}$ as compared to cells with an empty vector (pSEVA631). Dotted lines indicate induction with 1.0 µg mL$^{-1}$ aTc. Cell density ($OD_{600}$) (A) and median fluorescence intensity as analyzed by flow cytometry (B) are shown. The mean of biological quadruplicates is shown, with error bars representing the standard deviation.

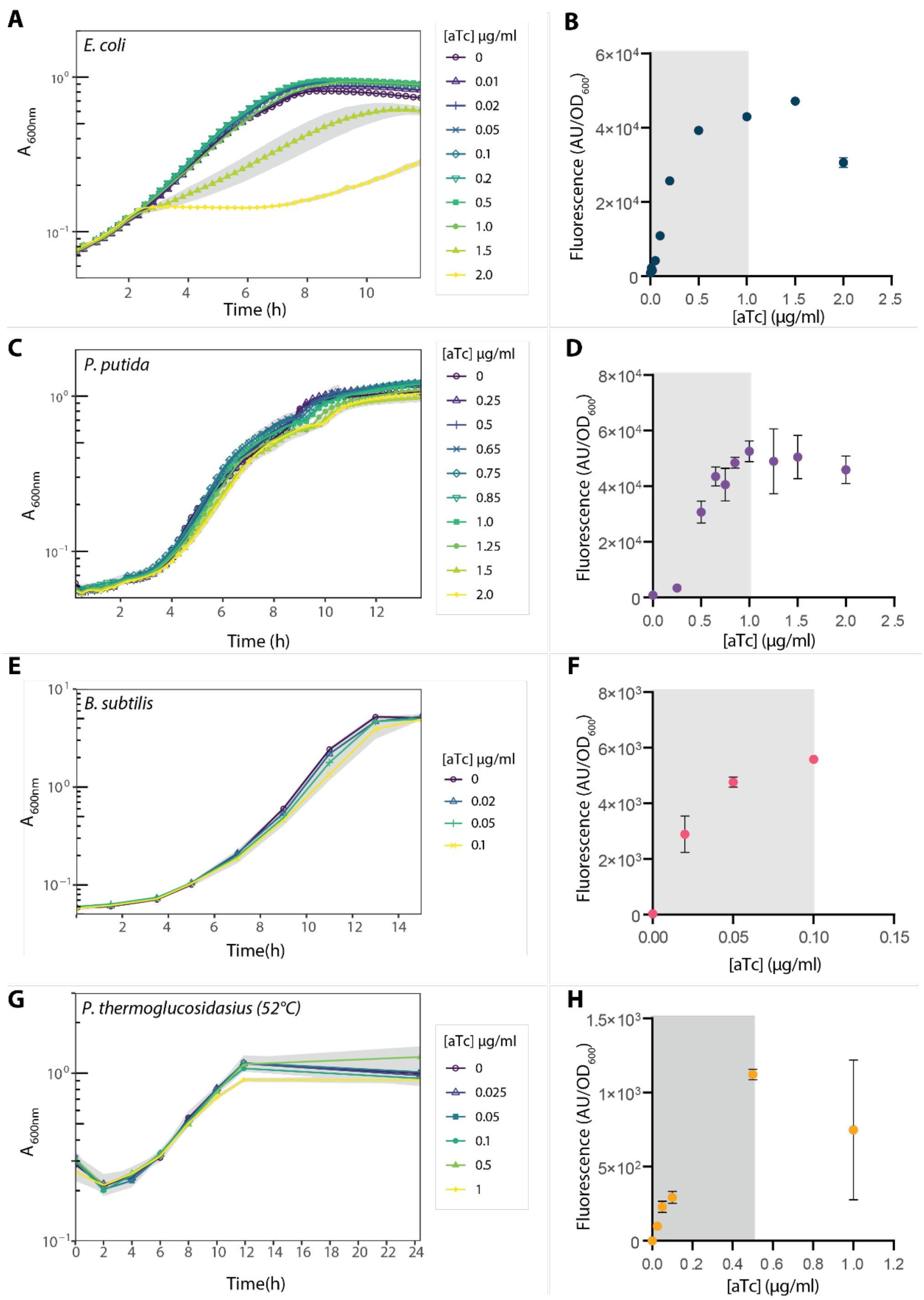


**Supplementary Figure S2**: Toxicity of aTc and titratability of TetR-$P_{tet}$* in *E. coli* (A, B), *P. putida* (C, D), *B. subtilis* (E, F), and *P. thermoglucosidasius* (G, H). Toxicity studies with various aTc concentrations were performed in 96-well microtiter plates, with $A_{600nm}$ values used as a proxy for $OD_{600nm}$. Previous work showed that *B. subtilis* is sensitive to aTc above 0.1 µg $mL^{-1}$, so higher concentrations were not tested here. Fluorescence was determined in a fluorescent microtitre plate reader and corrected for culture OD. All points represent the mean of biological triplicates, with standard deviation indicated in grey shading for toxicity or error bars for titratability data. The titratable range of tetR-$P_{tet}$* in each organism is shown through grey shading in B, D, F, and H.

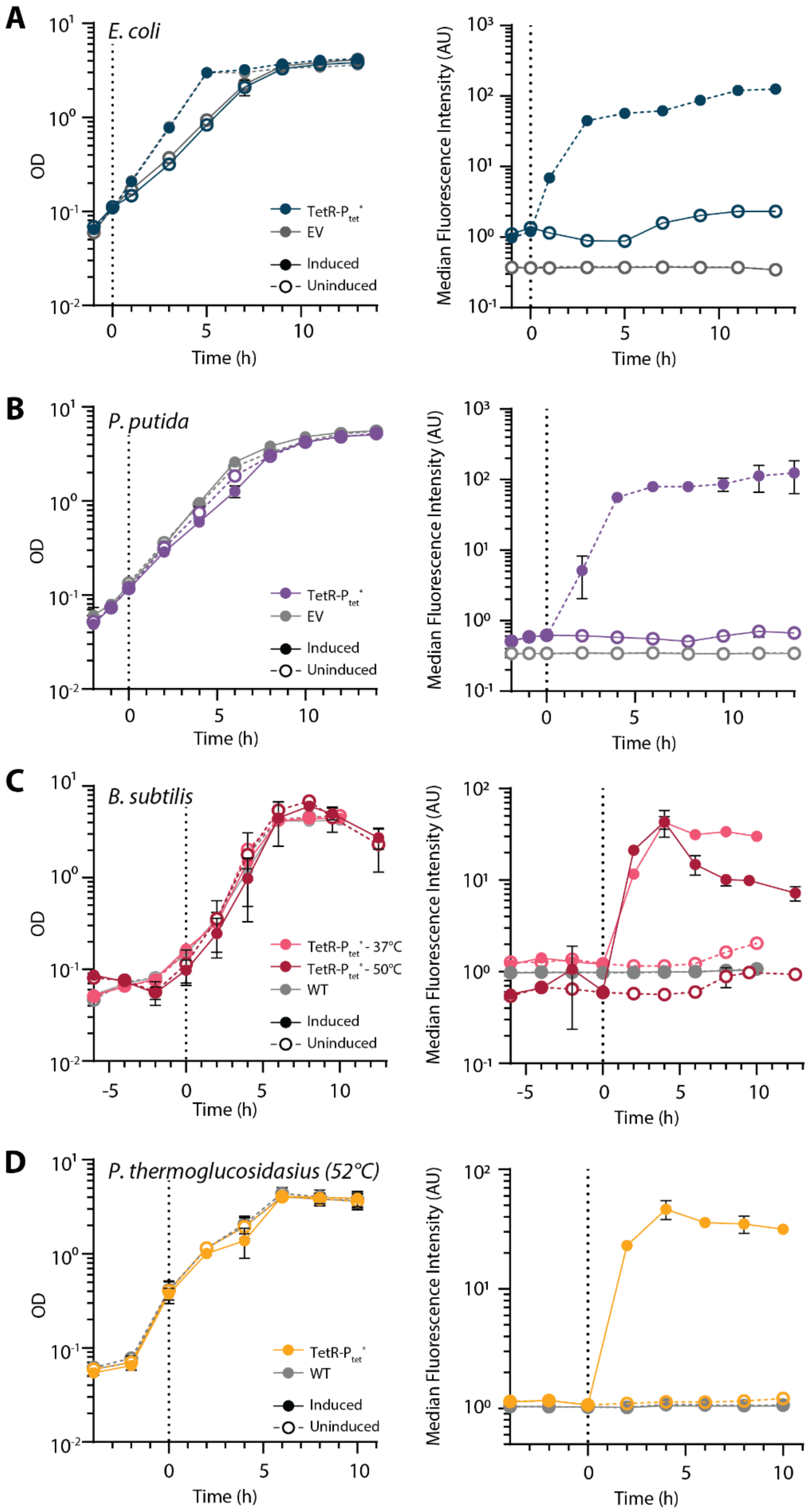


**Supplementary Figure S3:** Time course experiments of *E. coli* (A), *P. putida* (B), *B. subtilis* (C), and *P. thermoglucosidasius* at 52°C (D) expressing a tetR-$P_{tet}$* construct controlling mRuby2 expression. The left panels show measured cell density ($OD_{600}$), and the right panels show the median fluorescence intensity, as quantified by flow cytometry. Points indicate the mean of four biological replicates, with standard deviation depicted as error bars. The dotted line at $T_0$ indicates the time of induction.

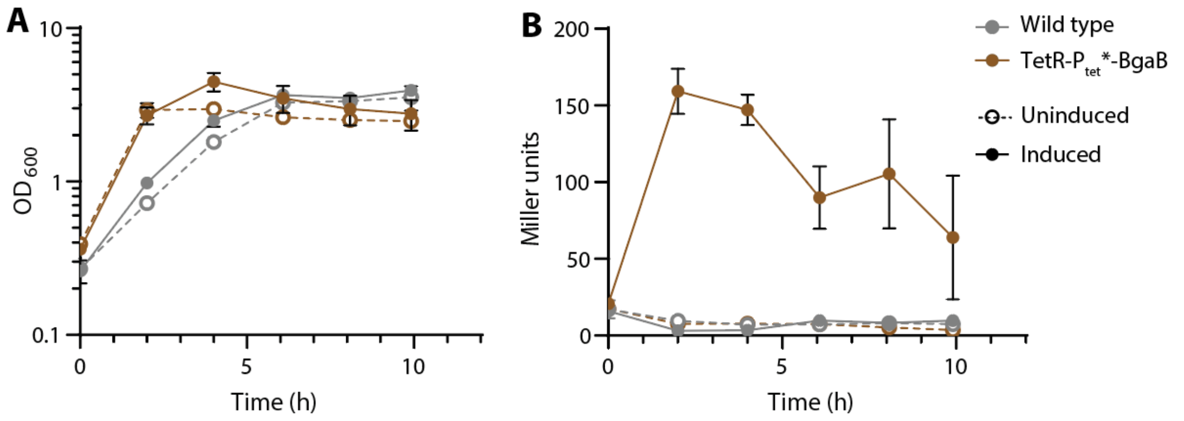


**Supplementary Figure S4:** Time course experiments of *P. thermoglucosidasius* expressing a tetR-$P_{tet}$* construct controlling BgaB expression at 60°C, with $OD_{600}$ (A) and BgaB activity expressed as miller units (B). Time indicates time post-induction and points indicate the mean of four biological replicates, with standard deviation depicted as error bars.

ProUSER X X X X X

| No. | Gram⁻ $Ab^R$ |
|---|---|
| 1 | Amp |
| 6 | Gm |
| 9 | $Kan^{Th}$ |

| No. | Gram⁻ OriV |
|---|---|
| 3 | pBBR1 |
| 4 | ColE1 |

| No. | Gram⁺ OriV |
|---|---|
| A | pHT315 |
| B | $pE194^{ts}$ |
| H | repB |
| I | pAMβ1 |
| **No.** | **Gram⁺ int. site** |
| C | *glmS* |
| D | *yrbD* |
| E | *yhgE* |
| F | *ΔsigF* |
| G | *rsbP* |
| J | repB-Δ*ldh*$^{Pt}$ |

| No. | Gram⁺ $Ab^R$ |
|---|---|
| 0 | Empty |
| 1 | $Cam^{BS}$ |
| 2 | $Ery^{BS}$ |
| 3 | $Spc^{BS}$ |
| 4 | $Kan^{BS}$ |

| No. | (Reg. &) Prom. |
|---|---|
| A | $P_{veg}$ |
| B | $P_{3P}$ |
| C | *manR* & $P_{manA}$ |
| D | $P_{mtlA}$ |
| E | *xylR* & $P_{xylA}$ |
| F | *tetR* & $P_{tet}$ |
| G | *tetR* & $P_{tet}$* |

**Supplementary Figure S5:** Overview of the updated ProUSER2.0 nomenclature to include the plasmids provided in this work[23]. Amp = ampicillin, Cam = chloramphenicol, Ery = erythromycin, Spc = spectinomycin, Kan = kanamycin. Superscript "BS" designates that the resistance marker is for B. subtilis. "Th" designates the thermostable variant of the Kanamycin resistance cassette.

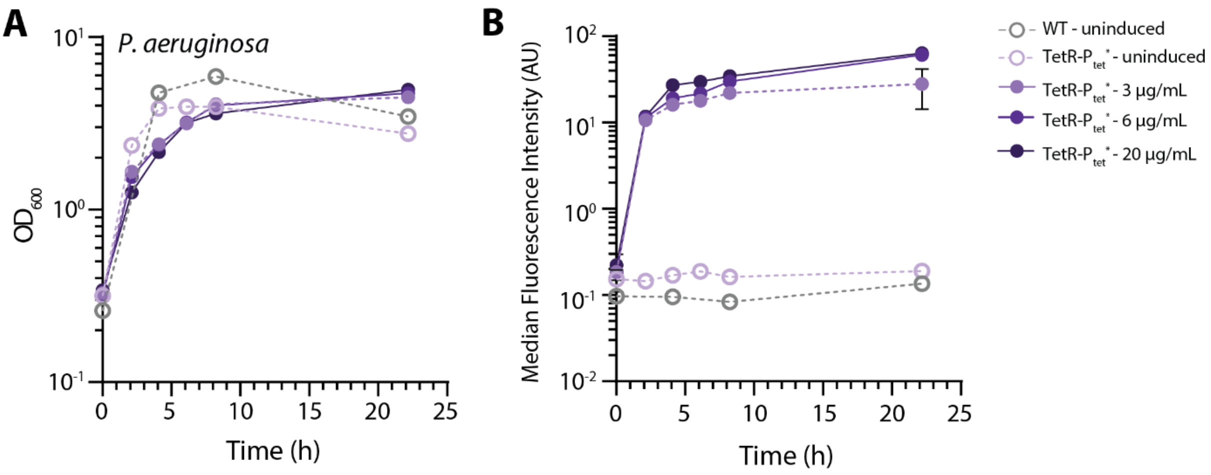


**Supplementary Figure S6:** Time course experiment of *P. aeruginosa* cultures expressing mRuby2 under the control of the TetR-$P_{tet}$* system when induced at different levels of aTc and a wild type, uninduced control. Dotted lines indicate no induction. Culture density ($OD_{600}$) (A) and median fluorescence intensity as analyzed by flow cytometry (B) are shown. The mean of biological quadruplicates is shown, with error bars representing the standard deviation.

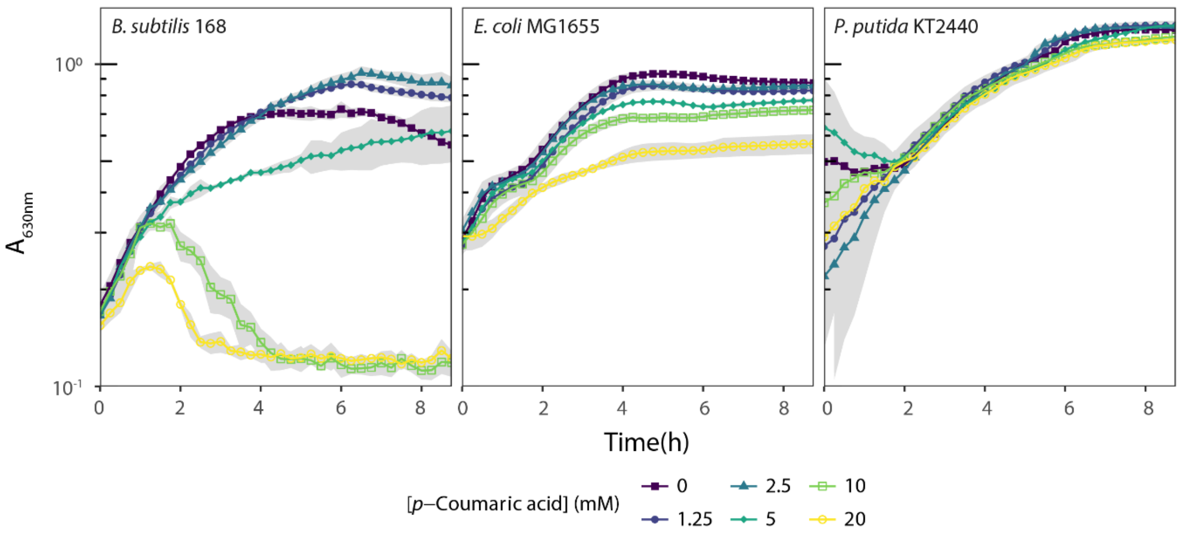


**Supplementary Figure S7:** Toxicity of *p*-coumaric acid in *B. subtilis*, *E. coli,* and *P. putida*. Absorbance was measured in a 96-microtiter well plate reader upon the addition of *p*-coumaric acid at the indicated final concentrations. Shaded areas indicate the standard deviation of biological triplicate measurements.

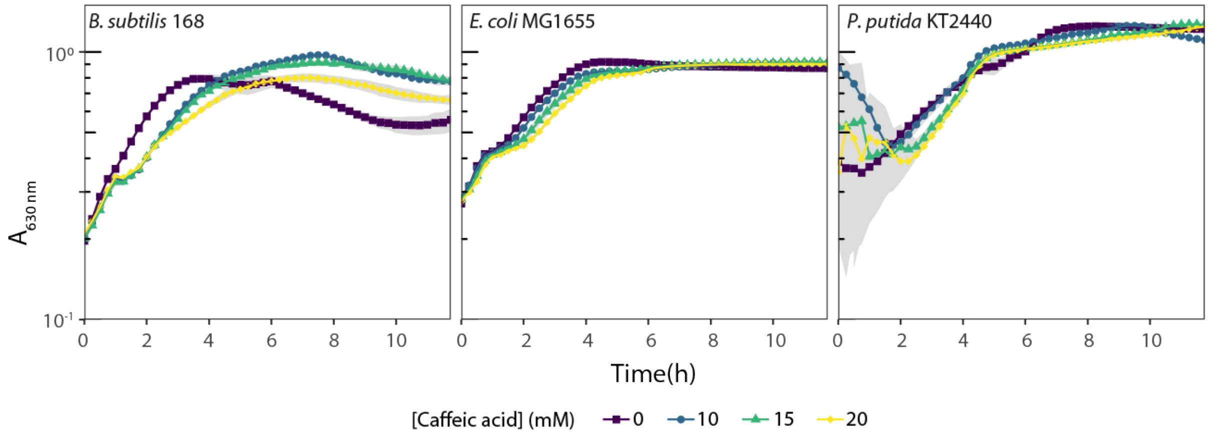


**Supplementary Figure S8:** Toxicity of caffeic acid in *B. subtilis*, *E. coli,* and *P. putida*. Absorbance was measured in a 96-microtiter well plate reader upon the addition of caffeic acid at the indicated final concentrations. Shaded areas indicate the standard deviation of biological triplicate measurements.

**Supplementary Table S1**: Overview of strains used in this study.

| **Strain** | **Genotype** | **Reference** |
|---|---|---|
| *E. coli* DH5α-λ***pir*** | F- φ80d*lacZ*ΔM15 Δ(*lacZYA-argF*)U169 *recA1 endA1 hsd*R17 (rk$^{-}$ mk$^{+}$) *sup*E44 *thi*-1 gyrA relA1 | [2] |
| *Escherichia coli* K-12 MG1655 | F$^{-}$ λ$^{-}$ *ilvG*$^{-}$ *rfb*-50 *rph*-1 | [3] |
| *Bacillus subtilis* 168 | *trpC2* | DSM23778 |
| *Pseudomonas putida* KT2440 | Mt$^{-2}$ *hsd*R1 (r$^{-}$ m$^{+}$) | [4] |
| *Parageobacillus thermoglucosidasius* DSM2542$^{T}$ | | DSM2542$^{T}$ |
| *Pseudomonas aeruginosa* PAO1 | | DSM1707 |
| SIJ650 | *B. subtilis 168,* glmS::$P_{mtlA}$-*comKS* | This work |
| SIJ590 | *B. subtilis* 168, glmS::TetR-$P_{tet}$-mRuby | [1] |
| VM47 | *P. thermoglucosidasius* Δldh:: TetR-$P_{tet}$*-mRuby | This work |
| VM57 | *P. thermoglucosidasius* Δldh:: TetR-$P_{tet}$*-BgaI | This work |
| SIJ678 | SIJ650 Δ*pad*::Spc$^{R}$ | This work |

**Supplementary Table S2**: List of plasmids used in this study. ProUSER2.0 indicates plasmids containing the proUSER2.0 cassette, as published previously[1]. "BS" designates that the integration site is for *B. subtilis*. Plasmid maps for all plasmids are provided as supplementary files.

| **Plasmid** | **Description** | **Reference** |
|---|---|---|
| pProUSER63H0G | BBR1-Gm$^{R}$-repB-Kan$^{R}$-TetR-$P_{tet}$* | This work |
| pProUSER63C1G | BBR1-Gm$^{R}$-glmS$^{BS}$-Cm$^{R}$-TetR-$P_{tet}$* | This work |
| pProUSER63I2G | BBR1-Gm$^{R}$-pAMβ1-Ery$^{R}$-TetR-$P_{tet}$* | This work |
| pProUSER94H0G | ColE1-Kan$^{R}$-repB-TetR-$P_{tet}$* | This work |
| pProUSER94J0G | ColE1-Kan$^{R}$-repB-Δ*ldh*-TetR-$P_{tet}$* | This work |
| pVM34 | ColE1-Kan$^{R}$-repB-Δldh-TetR-$P_{tet}$-mRuby | This work |
| pVM44 | BBR1-Gm$^{R}$-TetR-$P_{tetA(native)}$-mRuby | This work |
| pVM45 | BBR1-Gm$^{R}$-TetR-$P_{tet}$-mRuby | This work |
| pVM53 | ColE1-Kan$^{R}$-repB-Δldh-TetR-$P_{tet}$*-BgaI | This work |
| pVM61 | BBR1-Gm$^{R}$-glmS-Cm$^{R}$-TetR-$P_{tet}$-mRuby | This work |
| pVM62 | BBR1-Gm$^{R}$-pAMβ1-Ery$^{R}$-tetR-$P_{tet}$-mRuby | This work |
| pProUSER94H0G-mRuby2 | ColE1-Kan$^{R}$-repB-TetR-$P_{tet}$*-mRuby2 | This work |
| pProUSER63H0G-mRuby2 | BBR1-Gm$^{R}$-repB-Kan$^{R}$-TetR-$P_{tet}$*-mRuby2 | This work |
| pProUSER63H0G-C3H | BBR1-Gm$^{R}$-repB-Kan$^{R}$-TetR-$P_{tet}$*-C3H | This work |
| pProUSER63H0G-COMT | BBR1-Gm$^{R}$-repB-Kan$^{R}$-TetR-$P_{tet}$*-COMT | This work |
| pProUSER63H0G-synOMT | BBR1-Gm$^{R}$-repB-Kan$^{R}$-TetR-$P_{tet}$*-synOMT | This work |

**Supplementary Table S3:** Oligos used in this study

| No. | Name | Sequence (5' to 3') |
|---|---|---|
| P22 | Ptet_Fw_seq | GCGTTAACAGATCTGAGCTCCT |
| P23 | Ptet_FW_seq2 | TATTTCGATGCCCTGGACTT |
| P24 | mRuby_Rev_seq | CCATGTACGGATTGCCTTCT |
| P26 | KanR_seq_Rev | TCCGGATATTCTTCCAGCAC |
| P27 | glmsDS_seq_rev1 | AGCGTTGATCCAGGTACGTT |
| P31 | OriT_seq_FW | TGCTTCGGGGTCATTATAGC |
| P32 | OriT_seq_Rev | GGTGCGAATAAGGGACAGTG |
| P35 | bBR1_seq_rev | GGTTGGACACCAAGTGGAAG |
| P45 | kanRthermo_seq_rev | TGCCATGTTTCATTGCTCTC |
| P50 | LacZ_seq_Fw | TTTCGCTACCTGGAGAGACG |
| P55 | AmpR_seq_FW | TGGATGAACGTAACCGTCAG |
| P61 | BBR1_seq_FW | CTTCCACTTGGTGTCCAACC |
| P62 | RepB_seq_Fw | CTGAAAGGTGCGTTGAAGTG |
| P64 | mRuby_Unick_FW | GGCGAUAGGAGGAATATACATGGTGTCTAAGGGCGAAGAG |
| P73 | nicking_Rev | CAGTCACGACGCTGAGGTG |
| P77 | glmsUS_Rev | TAGACAGAAGCGGCATGTTG |
| P78 | glmsUS_FW | CTGAAAGGCCTAGACGATGC |
| P79 | CmRintFW | TGGTTACAATAGCGACGGAGA |
| P80 | glmsDS_FW | AAGGCAGCGCATTAGAACAT |
| P81 | BBR1_FW | AACAGCGATTCGTCCTGGT |
| P83 | KanR_Fw2 | ATCTGGCCATTCTGTGGAAC |
| P90 | AmpR_Rev | CGCTGTTCAGATCCAGTTCA |
| P108 | ldhR_Rev | GAATGCGGGAGTTAAACGAT |
| P109 | ldhP_seq | GTTGTAAAACGACGGCCAGTGC |
| P110 | ldhP_seq | GGCCGCTGTATCCATATGACCATG |
| P119 | ColE1_seq | AACCGTATTACCGCCTTTGA |
| P120 | ldhKO_seq | CTTAAACAAAAGCGTCGTTCGCTGC |
| P121 | ldhKO_seq | GATGAAATATTTCCCGTTTCAAGGGATCAGC |
| P124 | bgaB_U_Fw | GGCGAUAGGAGGAATATACCATGAACGTTTTATCCTCAATTTG |
| p125 | bgaB_U_Rev | AGAGTUAGAAAAAGAAACAGAGGCTACTCTCAA |
| P126 | UFW_OriT | AGTGCUGACCCTTGTCCTTTTCCGCTGCATAAC |
| P127 | URev_ColE1 | ATCGCCUCCTCTCAGGCGCCGCTGGTG |
| P128 | UFW_repB_kpn | ATCTGAAUGGTACCCTACTCTTTAATAAAATAATTTTTCCGTT |
| P129 | URev_repB_kfl | AGCACUGGGACCCCGAATTAATTCCTTAAGGAACGTACA |
| P130 | UFW_KanT | ATCGCACCUTCAAAATGGTATGCGTTTTGACACA |
| P131 | URev_KanT | ATTCAGAUCAGTTTGTTGAAGATTAGATGCTATAAT |
| P132 | UFW_tetR | ACGCAGUGGCGCGCCTGTACACAGCTGTCTAG |
| P135 | UFW_KanT_int | AGTGCUTCAAAATGGTATGCGTTTTGACACA |
| P136 | UFW_glmsUS_gth | ATCGCACCUTGTTAAGTGACAAAAAAGTCGTATACT |
| P138 | UFW_Gm | AGTGCUCAATAATTACGATTTAAATTTGACATAAGC |
| P139 | Urev_BBR1 | ATCGCCUCCCCCTACGGGCTTGCTCT |
| P140 | UFW_P15a | AGTGCUCAATAATTACGATTTAAATTTGTGTCTCAA |
| P141 | Urev_Km | ATCGCCUGGATATATTCCGCTTCCTCGCTC |
| P146 | glmsDS_fw | CAGAACGGGTATTTCTATTTTCG |
| P147 | glmsUS_rev | CCCCGCTTTGAGAAATGTAA |

| | | |
|---|---|---|
| P149 | Bgal_FW | CCCCTAGAAAGGCACCAACT |
| P150 | Bgal_Rev | CGGAAGTCCTGCATACCAAT |
| P151 | KanRth_Fw | TCACTTCCACCTTCCACTCA |
| P152 | ColE1_Rev | GGTTAACCCAAGAGCCCAAT |
| P278 | LDHDS_U_FW | ATCGCACCUTCAGTTAGGCACCGTGTATAAATGAAAAAAG |
| P287 | LDH_gFW | GCGTTTGGAGCGATAGGAAT |
| P312 | TER_U_Rev | AGGTGCGAUCTGGATTCTCACCAATAAAAAACG |
| P315 | dLDH_DS_Ufw | ATCGCACCUCCCTCGGCAAACAGAGCTTTA |
| P317 | BB_U_fw | AGTGCUCCTGCAGGGGGCCCT |
| P352 | VC9_BB_URev | AATTGUTATCCGCTTTAATTAAAGGCATCAA |
| P353 | tetRnew_UFW | ACAATUTCACACAGGAGGCCGA |
| P354 | TetRnew_URev | ATCGCCUCAGCTTACTGCAGGAGGAC |
| P355 | lacZ_UFW | AGGCGAUCGCAACGTCGTGACTGGGA |
| P356 | LacZ_Urev | AGCACUCTGGATTCTCACCAATAAAAAACG |
| P357 | colE1_FW | CAGCAGATTACGCGCAGAAA |
| P358 | colE1_FW2 | TTCCTCGCTCACTGACTCG |
| P490 | Tn7_Gm | ATATCGACCCAAGTACCGCC |
| P553 | pSEVA_seq_rv | CCGAGCGTTCTGAACAAATC |
| P587 | seq_seva_fw | TCTAGGGCGGCGGATTTG |
| P801 | Seva_ant_seq_fw | AAACCCTGGCGACTAGTCTT |
| P802 | Seva_ant_seq_rv | AGGAAAGTCTACACGAACCCT |
| P852 | Gth_BB_U_FW | AACAAGGUGAACATCGTGTGGATACAAC |
| P853 | Gth_BB_U_REV | AACTCUGCCATTATCATTTCCGTAATGCC |
| P1351 | lacZ_q_fw1 | TAATCGCCTTGCAGCACATC |
| P1352 | lacZ_q_rv1 | GACAGTATCGGCCTCAGGAA |
| P1353 | lacZ_q_fw2 | TTTCATCTGTGGTGCAACGG |
| P1354 | lacZ_q_rv2 | CGGTTTATGCAGCAACGAGA |
| P1431 | Seva_seq_fw | CGGCGGATTTGTCCTACTCA |
| P1567 | Seva_gadget_rv | GTAACATCGTTGCTGCTCCA |
| P1572 | tetR_seq_ekstra_fw | TCGCGATGACTTAGTAAAGCAC |
| P1573 | tetR_seq_fw | AAGCAGCTCTAATGCGCTGT |
| P1823 | CB_22_seq_tetR_rv | CGCCCAGAAGCTAGGTGTAG |
| P1824 | CB_23_seq_tetR_fw | TGCAGAGCCAGCCTTCTTAT |
| P1881 | Ruby_U_fw | GGCGAUAGGAGGAATACAATGGTGTCTAAGGGCGAAGA |
| P1882 | Ruby_U_rv | GGTGCGAUTTACTTGTACAGCTCGTCCATCC |
| P1928 | pAMB1_seq_fw6 | AGCGAAGCGAACACTTGATT |
| P1930 | pAMB1_seq_fw8 | TCCATGGACTTCATTTACTGG |
| P1949 | seva_seq_fw | TGTTCAGAACGCTCGGTTG |
| P1956 | HT315_seq_rv7 | CTGTTGTTTGTCGGTGAACG |
| P2072 | glmS_up_fw | CAACATGCCGCTTCTGTCTA |
| P2073 | glmS_down_rv | TTAGCCTTCGCGTACTCCTT |
| P2543 | Junk_seq_Fw | AGAGGGACTGCGACGTTCTA |
| P2546 | pGeo_seq_rev | ATTTTTCCGTTCCCAATTCC |
| P2561 | Rev_U_VC2_Ptet | ATCGCCUCAGCTTACTGCAGGAGCTCAGATCTG |
| P2564 | REV_U_VC2_Ter | AGGTGCGAUCCTGCAGGCTGGATTCTCACCAATAAAAAACGCCCGGC |
| P2585 | FW_U_VC_Junk_2 | AGGCGAUCTCGAGCTAGGTAACTAGAGGGA |
| P2617 | C3H_U_fw | GGCGAUAGGAGAAATACATATGACGATTACCTCTCCGGCA |

| | | |
|---|---|---|
| P2618 | C3H_U_rv | GGTGCGAUTTACGTGCCCGGGTTAATCA |
| P2619 | COMT_U_fw | GGCGAUAGGAGAAATACATATGGGCAGCACCGCAGAA |
| P2620 | COMT_U_rv | GGTGCGAUTTACAGCTTTTTCAGCAGTTCAATCA |
| P2621 | synOMT_U_fw | GGCGAUAGGAGAAATACATATGGGCAAAGGCATTACCGGT |
| P2622 | synOMT_U_rv | GGTGCGAUTCATTTTTTCAGTGCCAGGGTCA |
| P3065 | padC_up_KO_U | AGCTTGCAUATCGACCTCGCCGTTTATTGA |
| P3066 | padC_up_KO_U_rv | ACACGUTTACATCTTACACACTCTCCTTAGTCT |
| P3067 | padC_down_KO_U_fw | AGCACUTCGCGCGGGAAGATTATAA |
| P3068 | PadC_down_KO_U_rv | AGTCAGGUTCCAGAAAGCAATCATGGA |
| P3069 | PadC_check_fw | TTCTGCGAGCATTGTCTGTT |
| P3070 | padC_check_rv | TCTTCCTGTCATGGTGTGGA |

**Supplementary Table S4:** 16S sequences from corresponding strains used for generating the phylogenetic tree, indicated through NCBI accession numbers of the used sequences.

| Phyla | Strain | NCBI accession number |
|---|---|---|
| Actinobacteria | *Actinomyces oris* strain WVU 474 | NR_104896.1 |
| | *Corynebacterium glutamicum* MB001 | CP005959.1 |
| | *Mycobacterium tuberculosis* H37Rv | NR_102810.2 |
| | *Mycolicibacterium smegmatis* strain ATCC 19420 | NR_025311.1 |
| | *Streptomyces coelicolor* A3(2) | Y00411.1 |
| Bacteroidetes | *Prevotella intermedia* strain B422 | NR_026119.1 |
| | *Bacteroides fragilis* ATCC 25285T | X83935.1 |
| | *Bacteroides thetaiotaomicron* ATCC 29148 | L16489.1 |
| Chlamydiae | *Chlamydia pneumoniae* strain TW-183 | NR_026527.1 |
| | *Chlamydia trachomatis* strain HAR-13 | NR_025888.1 |
| Chlorobi | *Chlorobaculum tepidum* strain TLS | M58468.2 |
| | *Chlorobium chlorochromatii* strain CaD | NR_114906.1 |
| Cyanobacteria | *Prochlorococcus marinus* subsp. pastoris strain PCC 9511 | NR_125480.1 |
| | *Synechococcus elongatus* strain PCC7942 | CP000100.1 |
| | *Synechocystis* sp. PCC 6803 | KT371499.1 |
| Firmicutes (Bacilli) | *Bacillus amyloliquefaciens* DSM 7 strain ATCC 23350 | NR_118950.1 |
| | *Bacillus licheniformis* ATCC 14580 | X68416.1 |
| | *Bacillus subtilis* 168 | K00637.1 |
| | *Lactobacillus brevis* NS125 | EU177639.1 |
| | *Lactobacillus reuteri* C1 | EF412975.1 |
| | *Lactococcus lactis* subsp. lactis strain NCDO 604T | AB100803.1 |
| | *Parageobacillus thermoglucosidasius* DSM2542 | AB021197.1 |
| | *Staphylococcus aureus* ATCC 12600 | NR_118997.2 |
| Firmicutes (Clostridia) | *Acetobacterium woodii* DSM1030 | CP002987.1 |
| | *Caldicellulosiruptor bescii* DSM 6725T | CP001393.1 |
| | *Caldicellulosiruptor saccharolyticus* DSM 8903 | NR_074845.1 |
| | *Clostridioides difficile* 630 | NC_009089.1 |
| | *Clostridium acetobutylicum* ATCC 824 | NC_003030.1 |
| | *Clostridium ljungdahlii* DSM 13528 | FR733688.1 |
| | *Clostridium thermocellum* DSM 1313 | CP002416.1 |
| | *Thermoanaerobacterium thermosaccharolyticum* ATCC 7956 | FR870449.1 |

| Phyla | Strain | NCBI accession number |
|---|---|---|
| Fusobacteria | *Fusobacterium nucleatum strain* DNF00606 | KU726670.1 |
| | *Fusobacterium_necrophorum strain* RMA16505 | EF447427.1 |
| | *Ilyobacter insuetus* DSM 6831 | AJ307980.1 |
| | *Streptobacillus moniliformis* strain DSM 12112 | NR_027615.1 |
| Proteobacteria (Acidithiobacillia) | *Acidithiobacillus ferrooxidans* ATCC 23270 | AJ278719.1 |
| | *Acidithiobacillus thiooxidans* ATCC19377 | NR_115265.1 |
| Proteobacteria (Alpha) | *Bradyrhizobium japonicum* USDA122 | AF208503.1 |
| | *Methylobacterium extorquens* IAM 12631 | AB175632 |
| Proteobacteria (Beta) | *Cupriavidus necator* H16 | M32021.1 |
| | *Nitrosomonas europaea* C-31T (Nm50) | M96399.1 |
| | *Spirillum winogradskyi* D-427 | NR_115328.1 |
| Proteobacteria (Delta) | *Desulfovibrio salinus* P1BSR | NR_159913.1 |
| | *Desulfovibrio vulgaris* DSM 644 | AF418179.1 |
| | *Myxococcus xanthus* ATCC 25232 | NR_043945.1 |
| | *Sorangium cellulosum* DSM 14627 | NR_044443.1 |
| Proteobacteria (Epsilon) | *Campylobacter jejuni* NCTC 11168 | AL111168.1 |
| | *Nautilia profundicola* AmH | NR_041829.1 |
| | *Sulfurimonas autotrophica* OK10 | NR_028643.1 |
| | *Helicobacter pylori* ATCC 43504 | Z25741.1 |
| Proteobacteria (Gamma) | *Actinobacillus succinogenes* DSM 22257 | AF024525.1 |
| | *Cellvibrio japonicas* Ueda107 | NR_074804.1 |
| | *Escherichia coli* DH5a | CP026085.1 |
| | *Escherichia coli* K12 MG1655 | NR_102804.1 |
| | *Pseudomonas putida* KT2440 | AE015451.2 |
| | *Pseudomonas aeruginosa* DSM 50071 | CP012001.1 |
